# Adaptation to starvation requires a flexible circadian clockwork in *Neurospora crassa*

**DOI:** 10.1101/2022.06.06.494928

**Authors:** Anita Szőke, Orsolya Sárkány, Géza Schermann, Orsolya Kapuy, Axel C. R. Diernfellner, Michael Brunner, Norbert Gyöngyösi, Krisztina Káldi

## Abstract

The circadian clock governs rhythmic cellular functions by driving expression of a substantial fraction of the genome and thereby significantly contributes to the adaptation to changing environmental conditions. Using the circadian model organism *Neurospora crassa,* we show that molecular timekeeping is robust even under severe limitation of carbon sources, however, stoichiometry, phosphorylation and subcellular distribution of the key clock components display drastic alterations. Protein kinase A, protein phosphatase 2A and glycogen synthase kinase are involved in the molecular reorganization of the clock. RNA-seq analysis reveals that the transcriptomic response of metabolism to starvation is highly dependent on the positive clock component WC-1. Moreover, our molecular and phenotypic data indicate that a functional clock facilitates recovery from starvation. We suggest that the molecular clock is a flexible network that allows the organism to maintain a rhythmic physiology and preserve fitness even under long-term nutritional stress.

## Introduction

Circadian clocks are endogenous timekeeping systems allowing organisms to adapt to cyclic changes in the environment. Circadian clocks rely on transcriptional-translational feedback loops (TTFL) in which positive elements of the machinery activate the expression of oscillator proteins which in turn negatively feeds back on their own transcription. Circadian clocks are closely linked to metabolism. On the one hand, the clock rhythmically modulates many metabolic pathways, and on the other hand, nutrients and metabolic cues influence clock function. It is therefore not surprising that in human, conditions involving circadian rhythm dysfunction, such as shift work or jetlag, are associated with an increased risk of metabolic disorders including obesity, metabolic syndrome and type 2 diabetes (Baron and Reid, 2014).

Although the timing of nutrient availability affects the phase of the rhythm, the circadian oscillator can compensate for varying nutrient levels and maintain a constant period. Accurate synchronization of metabolic processes with recurrent environmental conditions, such as light-darkness or temperature fluctuations, may be particularly critical for adaptation to nutrient deprivation. Because glucose levels affect many signal transduction pathways as well as transcription and translation rates (Ashe et al., 2000; Corral-Ramos et al., 2021; Jona et al., 2000; Sancar et al., 2012), glucose deficiency may challenge the TTFL-based circadian clock to operate at a constant period. Mechanisms of nutrient compensation have been intensively investigated in the circadian model organism *Neurospora crassa*. In *Neurospora* the White-Collar-Complex (WCC) composed of the GATA-type transcription factors WC-1 and WC-2, and Frequency (FRQ) represent the core components of the circadian clock. The WCC supports expression of FRQ which then interacts with an RNA helicase (FRH) and the casein kinase 1a (CK1a) (Cheng et al., 2005; Gorl et al., 2001). The FRQ-FRH-CK1a complex facilitates phosphorylation and thus inhibition of the WCC resulting in rhythmic changes in WCC activity and *frq* levels (Larrondo et al., 2015; Querfurth et al., 2011; Schafmeier et al., 2006). During a circadian day FRQ is progressively phosphorylated, which reduces its inhibitory potential and leads to its degradation (Querfurth et al., 2011). The negative feedback loop is connected with a positive loop, in which FRQ supports the accumulation of both WC proteins (Cheng et al., 2001; Lee et al., 2000). Similar to other organisms, the *Neurospora* circadian clock supports rhythmic expression of about 10% of the genome (Hurley et al., 2014; Sancar et al., 2015). The WCC also acts as the major photoreceptor of *Neurospora* and transduces light information to the clock (Froehlich et al., 2002; He et al., 2002). In *Neurospora*, short-term (0-16 hr) glucose deprivation triggers compensatory mechanisms at the transcriptional and posttranscriptional levels that maintain expression levels of the core clock proteins, thereby keeping period length constant (Adhvaryu et al., 2016; Emerson et al., 2015; Gyongyosi et al., 2017; Olivares-Yanez et al., 2016).

Aim of this study was to characterize how chronic glucose deprivation affects the molecular clock and what role the circadian clock plays in the adaptation to starvation. We analyzed the transcriptome response to long-term glucose starvation in *wt* and the clock-less mutant *Δwc-1* and found that the WCC has a striking impact on nutrient-dependent expression of a large set of genes, including enzymes and regulators of carbohydrate, amino acid and fatty acid metabolism. We show that molecular timekeeping is robust even under severe limitation of carbon sources. Moreover, our data provide evidence that the TTFL is able to function in a wide range of stoichiometric conditions of its key elements, dependent on glucose availability. Our results show that *Neurospora* recovers faster from starvation in the presence of a functioning clock, suggesting a significant impact of the circadian clock on organismal fitness.

## Results

### Glucose-deprivation results in altered expression of core clock components

To assess how long-term glucose starvation affects clock function, liquid cultures of *wt Neurospora* grown at 2% glucose (standard medium) were transferred to a starvation medium with 0.01% glucose for 40 hours. Cultures were kept in constant light (LL) resulting in steady state levels of both *frq* RNA and protein. The growth of *Neurospora* virtually stopped after the medium change, while the expression of the major clock components changed characteristically (Figure 1A and B). Both WC-1 and WC-2 expression decreased gradually to about 15% and 20% of the initial levels, respectively (Figure S1A). The amount of FRQ remained relatively constant after glucose deprivation, but a mobility shift characteristic of hyperphosphorylation of the protein was observed (Figure 1B).

**Figure 1.**
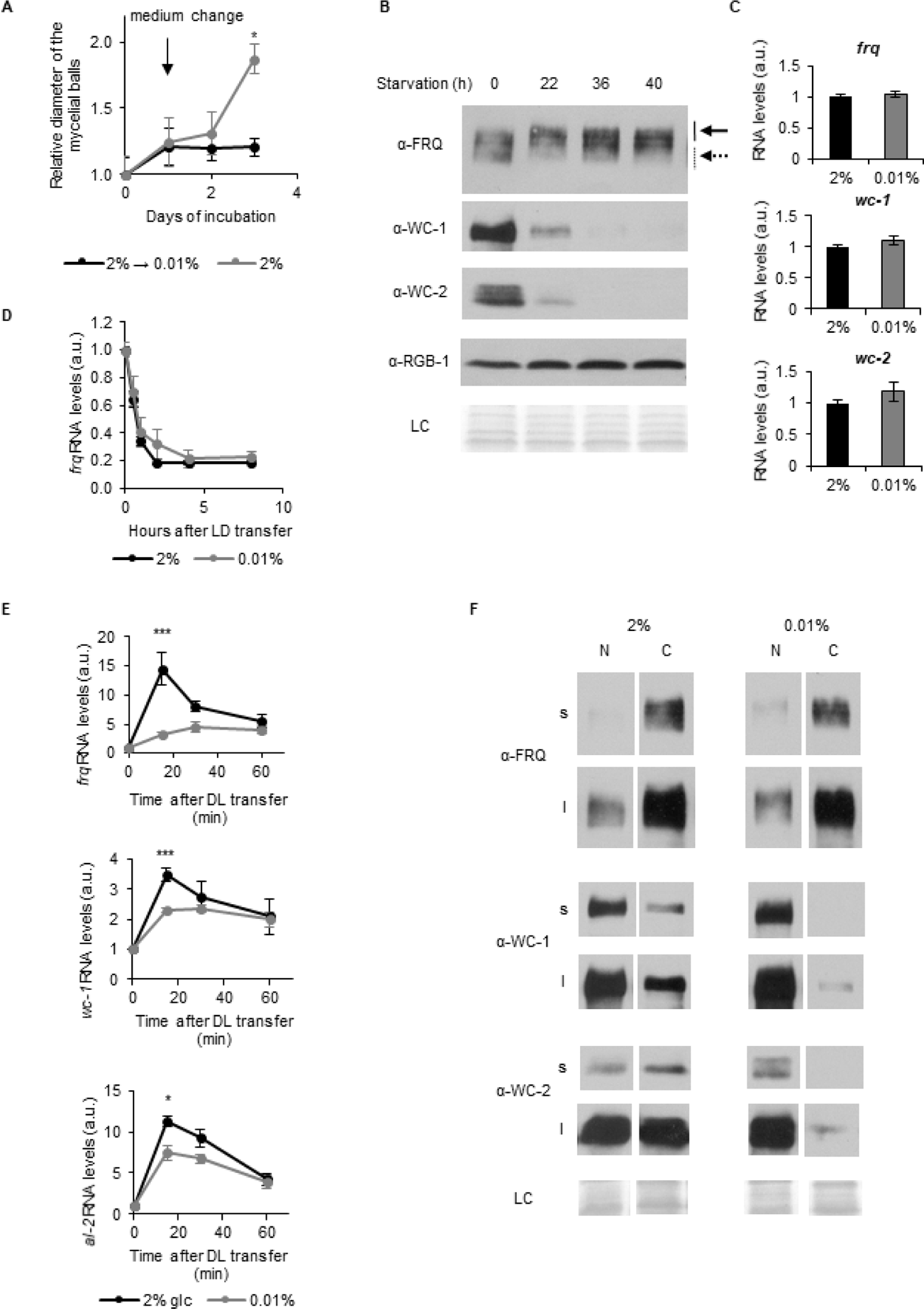
Glucose deprivation impacts both stoichiometry and subcellular distribution of clock components. (A) *Neurospora* growth is arrested in starvation medium. Following an incubation for 24 hours in standard liquid medium, mycelia were transferred to media containing either 2% or 0.01% glucose. Diameter of the mycelial balls was measured each day. The arrow indicates the time of medium change. (n=3, ±SEM, Repeated measures ANOVA, significant time-treatment interaction; post hoc analysis: Fisher LSD test) (B) Long term glucose starvation affects expression of clock proteins in *wt*. Mycelial discs were incubated for 24 hours in standard liquid medium and then transferred to starvation medium (time point 0). Samples were harvested at the indicated time points. Cell extracts were analyzed by Western blotting. Solid and dashed arrows indicate hyper- and hypophosphorylated forms of FRQ, respectively. RGB-1 and Ponceau staining (LC: loading control) are shown as loading controls. (n=3) See also Figure S1A. (C) RNA levels of *frq*, *wc-*1 and *wc-2* are similar under standard and nutrient limited conditions. Mycelial discs of the *wt* strain were incubated in standard liquid medium for 24 hours, then transferred to fresh media containing either 0.01% or 2% glucose and incubated for 40 hours in LL. RNA levels were normalized to that in cells grown in standard medium. (n=9-22, ±SEM, two-sample t-test, n.s.) (D) Stability of *frq* RNA is not affected by starvation. Growth conditions were described in (C). Following 40 hours of incubation in LL, cultures were transferred to DD (time point 0). Samples were harvested at the indicated time points. RNA levels were normalized to those measured at time point 0. (n=6, ±SEM, Repeated measures ANOVA, n.s.) (E) Light induction of gene expression is attenuated by glucose starvation. Mycelial discs of the *wt* strain were incubated in standard liquid medium for 24 hours, then transferred to media containing either 0.01% or 2% glucose. Following a 24-hour incubation in LL, cultures were transferred to DD for 16 hours and then light induced. Samples were harvested at the indicated time points after light on. Relative *al-2*, *frq* and *wc-1* RNA levels were normalized to that measured at the first time point. (n=5-11, ±SEM, Repeated measures ANOVA, significant time*treatment interaction, post hoc analysis: Tukey HSD test) (F) Glucose deprivation affects subcellular distribution of clock proteins. Growth conditions were as described in Figure 1C. Nuclear (N) and cytosolic (C) fractions were analyzed by Western blotting. (n=3, s: short exposure, l: long exposure, LC: loading control)

Because hyperphosphorylated FRQ exerts a reduced negative feedback on WCC (Schafmeier et al., 2006), we hypothesized that the starvation-induced phosphorylation of FRQ might lead to an increase in WCC activity and consequently to an acceleration of its decay (Kodadek et al., 2006; Punga et al., 2006; Schafmeier et al., 2008). To assess WCC stability, we followed WC levels in cultures treated with the translation inhibitor cycloheximide (Figure S1B). Our data suggest that the increased turnover of WC-1 may be partly responsible for the low WCC levels in the starved cells. In LL WCC promotes transcription of *frq* and *wc-1*. Although WCC levels were significantly different under standard and glucose-starved conditions, RNA levels of *frq* and *wc-1* were similar (Figure 1C), suggesting a compensatory mechanism that either maintains the active pool of WCC under various nutritional conditions constant or stabilizes the RNA. We examined whether glucose-deprivation affects stability of *frq* RNA. After light-dark transfer (LD), transcription of *frq* is repressed, and therefore changes in *frq* levels reflect RNA degradation. However, *frq* RNA levels after LD transition were similar under both culture conditions, suggesting that changes in RNA stability do not contribute to the maintenance of *frq* levels during starvation (Figure 1D). RNA levels of *wc-2* were also similar in starved and control cells, but expression of *wc-2* is not controlled by the WCC (Figure 1C).

To further examine the activity of the WCC, we investigated the expression of the light-inducible genes *frq*, *wc-1* and *al-2* after dark-light (DL) transfer under both nutrient conditions. The light-induced initial increase of RNA levels was lower in starved than non-starved cultures, whereas the steady state expression levels after light adaptation were similar under both conditions (Figure 1E). The difference in the kinetics of light induction suggests that the light-inducible pool or the photoreceptor function of WCC is reduced upon glucose deprivation.

Nucleocytoplasmic distribution of clock proteins is tightly associated with their phosphorylation and activity. Hence, we performed subcellular fractionation on our LL samples (Figure 1F). In accordance with previous data (Cheng et al., 2005; Gyongyosi et al., 2017; Schafmeier et al., 2006), the majority of FRQ was in the cytosol fraction, and its distribution did not change markedly upon glucose-deprivation. In contrast, WC proteins were virtually absent from the cytosol of starved cells, whereas their nuclear concentrations were similar to those in the control cells.

Although the expression of FRQ and WCC is interdependent (Cheng et al., 2001), their levels did not change proportionally upon starvation, raising the question of whether the oscillator function is intact under starvation conditions. We followed clock protein levels in constant darkness (DD), when the circadian clock displays a free running endogenous rhythm. FRQ protein showed a similar robust oscillation in both standard and starvation media, with no noticeable difference in period or phase (Figure 2A). When protein samples were analyzed on the same gel, increased FRQ phosphorylation was observed under starvation conditions in LL and at all time points in DD (Figure 2B). Similar to the changes in LL, the expression of WC proteins was greatly reduced in DD upon glucose deprivation. However, neither phase nor amplitude of *frq* RNA oscillation was affected by starvation (Figure 2C, left panel), indicating that WCC activity was similar under both conditions. Since starvation does not affect *frq* RNA decay (see above), an unknown mechanism must recalibrate the central clockwork to keep *frq* transcript levels and oscillation glucose-compensated despite the decline in WCC levels. To examine clock output function, we measured RNA levels of two clock-controlled genes, *ccg-2* and *fluffy* (Bell-Pedersen et al., 1992). Starvation resulted in significant upregulation of *ccg-2* expression similar to previous findings (Bell-Pedersen et al., 1992; Kaldenhoff and Russo, 1993; Sokolovsky et al., 1992). In addition, a robust oscillation of *ccg-2* and *fluffy* RNA was detected under both conditions, with peaks and troughs at the expected circadian time (Figure 2C, right panel; Figure S2).

**Figure 2.**
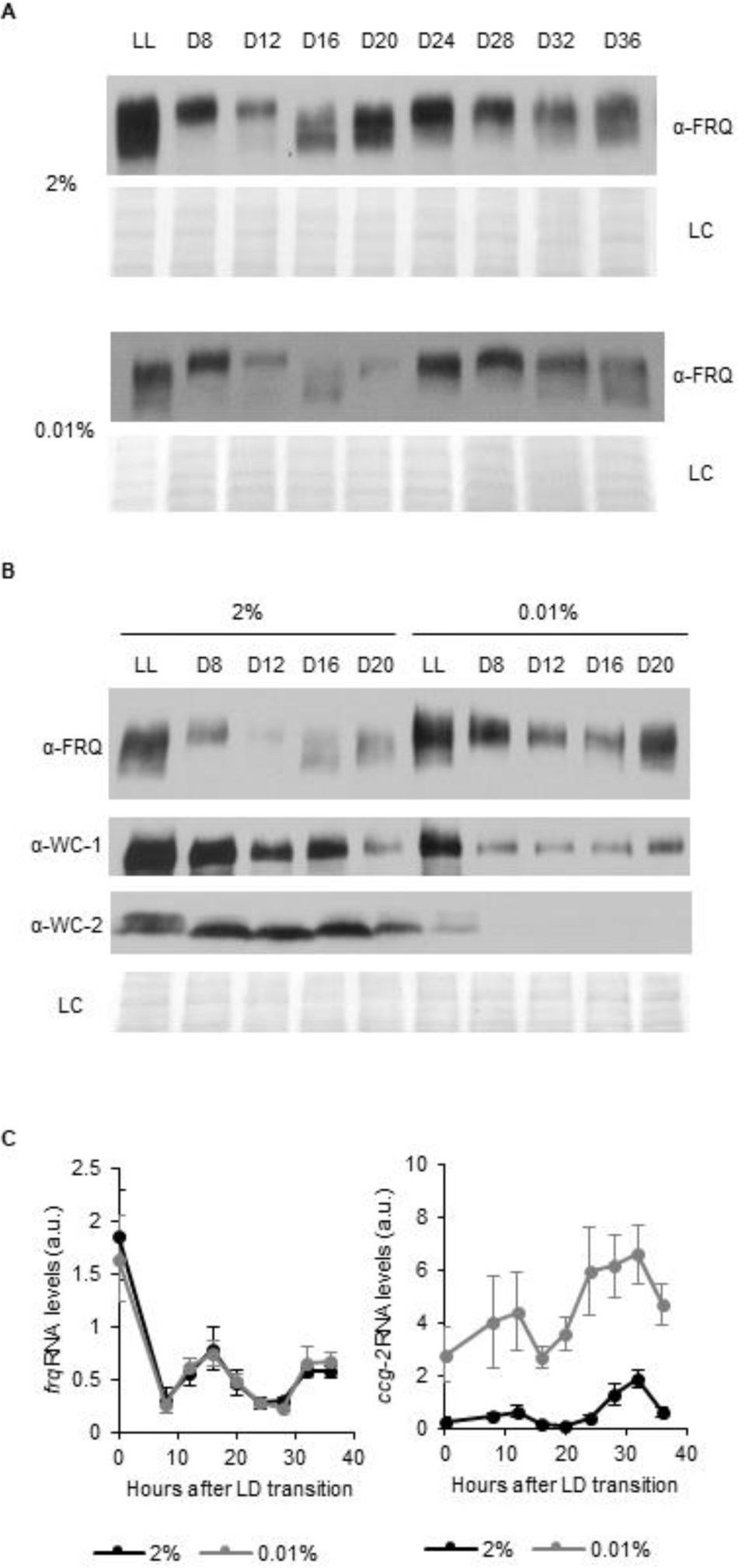
Effect of starvation on the molecular clock under free running conditions. (A) FRQ level oscillates under starvation conditions in DD. Following an incubation in standard liquid medium for 24 hours, mycelia were transferred to standard or starvation medium. After 24-hour incubation in LL, cultures were transferred to DD. Samples were harvested at the indicated time points. (n=3, LC: loading control) (B) WC levels are reduced and FRQ is hyperphosphorylated during glucose starvation. The experiment was performed as described in (A). Cell extracts from both growth conditions were analyzed on the same gel. (n=3, LC: loading control) (C) Expression of *frq* and *ccg-2* is rhythmic during long-term glucose starvation. Experiment was performed as described in (A). Relative RNA levels were determined. (n=3-11, ±SEM, Repeated measures ANOVA, n.s.)

Our results suggest that the circadian clock functions robustly during glucose deprivation despite increased FRQ phosphorylation and decreased WCC levels and drives rhythmic expression of output genes without changes in period length or phase.

### Multiple modulators are involved in the starvation response of the circadian oscillator

To address the role of FRQ-mediated feedback in the starvation response, we used the FRQ-less mutant *frq*^9^. In *frq*^9^, due to a premature stop codon, only a truncated, non-functional version of FRQ is expressed, but activity of the *frq* promoter can be determined by measuring *frq*^9^ RNA levels (Aronson et al., 1994; Liu et al., 2019). Because of the lack of the positive feedback of FRQ on the accumulation of the WC proteins, both WC-1 and WC-2 levels are low in *frq*^9^ (Cheng et al., 2001; Schafmeier et al., 2006). As shown in Figure 3A, absence of a functional FRQ protein attenuated the effects of starvation, i.e. amount of WC-1 did not change in response to glucose deprivation, whereas WC-2 levels were moderately reduced. Consistent with previous observations (Schafmeier et al., 2005), the increased expression of *frq^9^* in standard medium compared with *frq*, can be attributed to the absence of negative feedback from FRQ to WCC. However, while starvation did not alter *frq* levels in *wt*, it decreased the amount of *frq*^9^ RNA (Figure 3B). This suggests that FRQ contributes to both WCC depletion and FRQ level compensation. Action of CK1a on the clock proteins is dependent on its binding to FRQ via FRQ-CK1a-interaction domain 1 (FCD1) (Querfurth et al., 2011) and FCD2 (He et al., 2006). To examine the glucose-dependent impact of CK1a on the circadian clock, we used a *frqΔFCD1-2* strain. FRQΔFCD1-2 cannot recruit CK1a, and hence, the CK1a-dependent phosphorylation and inactivation of the WCC are impaired (Querfurth et al., 2011). Upon glucose withdrawal FRQΔFCD1-2 also displayed an electrophoretic mobility shift, and levels of both WC-1 and WC-2 were reduced, however, extent of the change was moderate compared to *wt* (Figure 3C, upper panel). *frqΔFCD1-2* RNA levels were low under standard conditions and did not change upon glucose deprivation (Figure 3C, lower panel). Our data suggest that stable recruitment of CK1a to FRQ is not essential for starvation-dependent hyperphosphorylation of FRQ but it affects WC levels. Both FRQ and WC-1 can be modified by PKA. Moreover, PKA activity decreases in both *Neurospora* and yeast when glucose is limited (Conrad et al., 2014; Li and Borkovich, 2006b). In the *mcb* strain the regulatory subunit of PKA is defective and hence, PKA is constitutively active at an elevated level. In accordance with literature data, FRQ protein was elevated in *mcb* (Huang et al., 2007) (Figure 3D). However, glucose deprivation did not cause a reduction of WC-1 or WC-2 levels and *frq* RNA became elevated. This indicates that PKA plays a prominent role in transducing starvation signals to the circadian clock.

**Figure 3.**
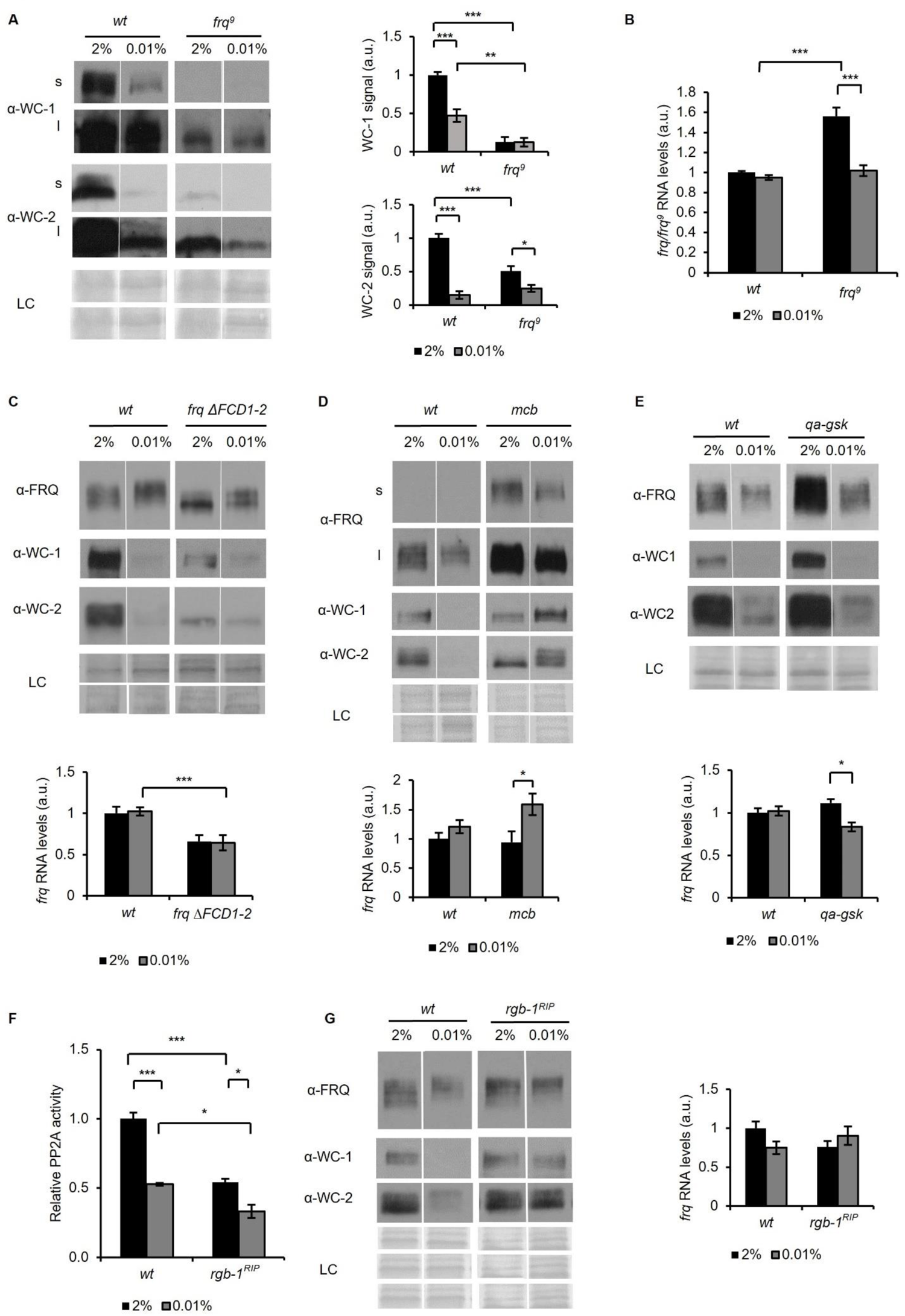
FRQ, PKA, GSK and PP2A affect the starvation response of the Neurospora clock. (A) Effect of starvation on WC levels is reduced in *frq*^9^. Growth conditions were as described in Figure 1C. Cell extracts were analyzed by Western blotting (left panel). (n=3) Protein signal density was analyzed (right panel). (n=3, ±SEM, LC: loading control for WC2 (upper panel) and WC1 (lower panel), Factorial ANOVA; significant strain*treatment interaction; post hoc analysis: Tukey HSD test) (B) *frq*^9^ RNA expression is sensitive to glucose deprivation. Growth conditions were as described in Figure 1C. RNA levels were normalized to that of *wt* grown in standard medium. (n=6, ±SEM, Factorial ANOVA; significant strain*treatment interaction; post hoc analysis: Tukey HSD test) (C) Impaired FRQ-CK1a interaction affects the starvation response of the molecular clock. Experiments were performed with the indicated strains as described in Figure 1C. Indicated protein (upper panel) and *frq* expressions (lower panel) were analyzed. (n (protein analysis) =12, LC: loading control for FRQ and WC2 (upper panel) and WC1 (lower panel), n (RNA analysis) =4-5, ±SEM, Factorial ANOVA; significant strain effect; post hoc analysis: Tukey Unequal N HSD test) (D) The starvation response is altered in the PKA mutant (*mcb*). Experiments were performed with the indicated strains as described in Figure 1C. Left panel: analysis of cell extracts by Western blotting (n=12, s: short exposure, l: long exposure; LC: loading control for FRQ (upper panel), WC1 and WC2 (lower panel)) Right panel: *frq* RNA levels of the indicated strains. (n=8-9, ±SEM, Factorial ANOVA; significant treatment effect; post hoc analysis: Tukey Unequal N HSD test) (E) Hyperphosphorylation of FRQ upon glucose withdrawal is dependent on GSK. Experiments were performed with the indicated strains as described in Figure 1C. The medium was supplemented with 1.5*10^-5^M quinic acid (QA) during the first day of incubation. Following the medium change, mycelia were incubated in QA-free medium. Upper panel: cell extracts analyzed by Western blotting. (LC: loading control) Lower panel: *frq* RNA levels of the indicated strains. (n=6, ±SEM; Factorial ANOVA, significant strain*treatment interaction, post hoc analysis: Tukey HSD test) (F) PP2A activity is decreased under starvation conditions. Experiments were performed with the indicated strains as described in Figure 1C. PP2A-specific activity of the cell lysates was determined and normalized to that of the *wt* grown in standard medium. (n=3-4, ±SEM, Factorial ANOVA, Significant strain*treatment interaction, post hoc analysis: Tukey Unequal N HSD test) (G) The starvation response is altered in the strain lacking a functional PP2A regulatory subunit (*rgb*-*1*^RIP^). Experimental procedures were performed with the indicated strains as described in Figure 1C. Cell extracts were analyzed by Western blotting (n=12, LC: loading control for FRQ (upper panel), for WC1 (middle panel) for WC2 (lower panel)) (left panel) and RNA levels of *frq* were determined. (n=9-10, ±SEM, Factorial ANOVA, significant strain*treatment interaction) (right panel)

Glycogen synthase kinase (GSK) is an important factor of the starvation response in many organisms including yeast (Quan et al., 2015) and was shown to fine-tune the circadian period in *Neurospora crassa* (Tataroglu et al., 2012). As GSK is an essential protein, we used the *qa-gsk* strain in which GSK expression is under the control of the quinic acid (QA)-inducible *qa-2* promoter (Tataroglu et al., 2012). Similar to previous findings (Tataroglu et al., 2012), without QA but in the presence of 2% glucose WC-1 and FRQ levels were elevated in *qa-gsk* (Figure 3E). The starvation response of the clock was partially affected by GSK depletion (Figure 3E). While WC levels changed similarly to those in *wt*, FRQ did not become hyperphosphorylated and *frq* RNA levels moderately decreased upon starvation. These results suggest that GSK supports the hyperphosphorylation of FRQ during starvation, but this modulation of FRQ is not sufficient to impact WCC levels.

PP2A is a major regulator of the clock affecting both FRQ and WCC (Colot et al., 2006; Gyongyosi and Kaldi, 2014; Querfurth et al., 2011; Schafmeier et al., 2006) and its activity is glucose-dependent in yeast (Hallett et al., 2014). As shown in Figure 3F, starvation leads to a decrease in PP2A activity in *Neurospora* to levels characteristic for the *rgb-1^RIP^* mutant, which lacks one of the regulatory subunits of the phosphatase complex. We have previously shown that dephosphorylation of WC-2 in CHX-treated cells depends on PP2A (Gyongyosi et al., 2013; Schafmeier et al., 2005). Hence, the delayed dephosphorylation of WC-2 in the starved cells suggest that glucose deprivation reduces WCC-modifying PP2A activity (Figure S1B). Next, we examined the effect of glucose-starvation on the clock in *rgb-1*^RIP^. While *frq* RNA and FRQ protein levels were similar in *wt* and *rgb-1*^RIP^, WC protein levels were not reduced in starved *rgb-1*^RIP^ (Figure 3G), suggesting that PP2A is involved in the glucose-dependent reorganization of the clock.

### Glucose-deprivation differentially impacts the transcriptome in *wt* and *Δwc-1*

To examine the role of the WCC in the adaptation to glucose starvation, we performed RNA-seq analysis in *wt* and *Δwc-1*. Liquid cultures were grown under standard and starving conditions in 12-12-hour dark-light cycles and samples were harvested after 48 hours (ZT 12, end of the light period). Under these conditions the clock is entrained in *wt*, and at ZT 12 an adapted state for light-dependent genes can be expected. Long-term glucose starvation affected more than the 20 % of coding genes at the level of RNA. The most upregulated genes in *wt* encoded polysaccharide degrading enzymes, conidiation specific proteins and monosaccharide transporters, similarly to findings in *Aspergillus niger* (Nitsche et al., 2012). Polysaccharide degrading enzymes (xylanases, endoglucanase) are considered scouting factors and their induction might play a foraging role increasing survival chances under starvation conditions (Benocci et al., 2017). Importantly, in *Δwc-1* 11 of the 15 most upregulated genes responded with a weaker increase to glucose-deprivation, indicating a limited transcriptomic adaptation when compared with *wt* (Table 1).

**TABLE 1.** The first 15 most upregulated annotated genes in *wt* during glucose starvation. For experimental procedures of the RNA-seq analysis see STAR Methods section. Genes with more pronounced upregulation in *wt* compared to *Δwc-1* are bold-lettered.

| ID | Name | Upregulation<br>in *wt*<br>(fold change) | Upregulation<br>in $\Delta wc-1$<br>(fold change) | *wt*/$\Delta wc-1$<br>(2%) | Gene product |
| --- | --- | --- | --- | --- | --- |
| NCU02500 | *ccg-4* | 281 | n.s. | n.s. | clock-controlled pheromone |
| NCU07225 | *gh11-2* | 254 | 163 | n.s. | xylanase |
| NCU08769 | *con-6* | 117 | 88 | n.s. | conidiation specific protein |
| NCU05924 | *gh10-1* | 96 | 15 | n.s. | xylanase |
| NCU00943 | *tre-1* | 77 | 102 | n.s. | trehalase |
| NCU07325 | *con-10* | 70 | 13 | n.s. | conidiation specific protein |
| NCU08189 | *gh10-2* | 67 | 13 | n.s. | xylanase |
| NCU10055 | *nop-1* | 66 | 58 | n.s. | opsin |
| NCU06905 | *thnr* | 65 | n.s. | 0.01 | tetrahydroxynaphthalene<br>reductase |
| NCU08457 | *eas* | 49 | 73 | n.s. | hydrophobin |
| NCU09873 | *acu-6* | 39 | 12 | 2.9 | phosphoenolpyruvate<br>carboxykinase |
| NCU10021 | *hgt-1* | 35 | 56 | 1.4 | monosacharide transporter |
| NCU08755 | *gh3-3* | 28 | 38 | n.s. | beta-glucosidase |
| NCU00762 | *gh5-1* | 21 | n.s. | n.s. | endoglucanase |
| NCU08114 | *cdt-2* | 21 | 11 | n.s. | hexose transporter |

In further analyses we aimed to identify genes that responded differently to glucose withdrawal in *wt* and *Δwc-*1 or showed differences in their *Δwc-1/wt* expression ratios between starved and non-starved conditions. We set a two-fold significant difference as threshold for physiologically relevant changes (Figure 4A, B). The detailed analysis showed that a sum of 1377 RNAs representing 13% of the 9758 coding genes changed in a strain-specific manner in response to glucose starvation (Figure S3). Detailed comparison of the *Δwc-1/wt* ratios of transcript levels revealed that 1348 genes were differently expressed in the two strains at either standard or starved conditions (Figure S4).

**Figure 4.**
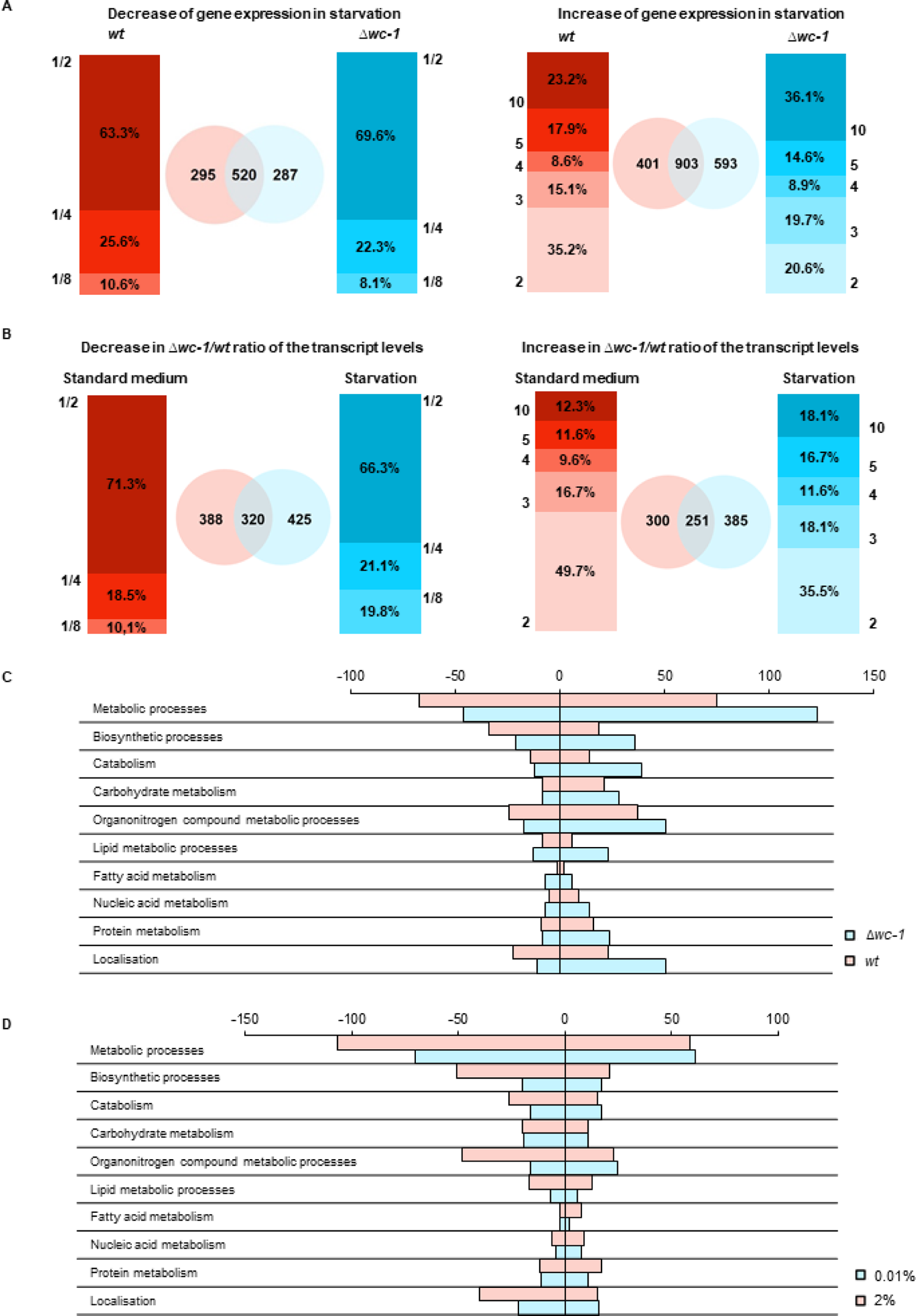
WC-1 is required for adaptation to starvation in genome-wide scale. (A) Distribution of number of genes showing starvation induced up- and downregulation in *wt* and *Δwc-1*. Values on the y-axis of the bar graphs indicate the minimal and maximal fold-change of up- and downregulation, respectively. Venn-diagrams indicate the number of up- and downregulated genes in *wt* (red) and *Δwc-1*(blue). (B) Distribution of genes expressed at lower and higher level in *Δwc-1* than in *wt* in standard or starvation medium. Values on the y-axis of the bar graphs indicate the minimal and maximal ratios of RNA levels (*Δwc-1*/*wt*). Venn-diagrams indicate the number of genes showing different expression in the two strains in the indicated medium. (red: standard medium, blue: starvation) (C) Number of genes showing strain-specific changes in major metabolic functions in response to a 40-hour glucose deprivation. Positive and negative values indicate number of genes with increased and decreased RNA levels, respectively. Genes were classified by GO analysis (Mi et al., 2013). (D) Number of genes showing treatment-specific (2% vs 0.01% glucose) changes in their *Δwc-1*/*wt* RNA ratio. Positive and negative values indicate number of genes with increased and decreased RNA ratio, respectively. Genes involved in major metabolic functions were classified by GO analysis (Mi et al., 2013).

Using the Gene Ontology (GO) enrichment tool (Mi et al., 2013), we found that a substantial proportion of the genes responding differently to starvation in *wt* and *Δwc-1* is involved in crucial metabolic processes (Figure 4C, D, Figure S5, Table S1). Further analysis showed that in the *wt* the downregulation of biosynthetic processes of small molecules and amino acids dominated, whereas *Δwc-1* had a higher tendency to up-regulate amino acid catabolism and down-regulate fatty acid synthesis in response to long-term starvation (Table S2). In analysis using the KEGG database, strain-specific differences in genes assigned to glycolysis, citrate and glyoxylate cycle, the pentose phosphate pathway and amino acid metabolism were identified (Figure S6-S8, and Table S3). However, pathways of fatty acid biosynthesis displayed the most uniform strain-dependent differences in response to starvation with a dominant downregulation in Δ*wc-1* compared to *wt* (Figure S8 and Table S3). Among constituents of these pathways only tetrahydroxynaphthalene reductase-2 was strain-specifically upregulated in *wt* (Table 1). Importantly, this enzyme catalyzes key steps of the synthesis of melanin, which is a crucial factor required for mechanical strength of cell wall and is therefore central in the adaptation to extreme changes in the environmental conditions (Nosanchuk et al., 2015).

To validate RNA-seq data, we performed qRT-PCR measurements for genes involved in the metabolism of carbohydrates or amino acids as well as for genes important for conidiation, the well-characterized nutrient-dependent process in fungi (Springer, 1993) (Figure S9). All tested RNAs showed similar expression ratios in both analyses indicating the high reliability of our transcriptome profiling data (Table S4). Striking differences between *Δwc-1* and *wt* were observed for the conidiation-specific protein 10 (*con-10*, NCU07325), the conserved mitochondrial gene *tca-3* (NCU02366), encoding a putative aconitase that catalyzes the citrate-isocitrate transition, and the gene of choline dehydrogenase (*choldh, NCU01853*) promoting the production of glycin-betain from choline, and thereby participating in amino acid metabolism.

### Efficient recovery from starvation is dependent on a functional clock

Based on the transcriptomic data indicating significant differences between *wt* and *Δwc-1* in the adaptation to glucose limitation, we hypothesized that starved *wt* might be better prepared to growth regeneration and could respond more effectively to the resupply of glucose than clock deficient strains. *wt* and the clock mutants *frq*^10^ and *Δwc-1* were incubated in light-dark cycles and following the 40 hours starvation the medium was supplemented with 2% glucose (Figure 5A). Interestingly, the medium of starved *wt* cultures was cloudy before the addition of glucose, whereas the medium of clock-less cultures remained clear (Figure 5B, Before: upper panel). As observed after filtration of the cultures (Figure 5B, Before: lower panel), the cloudy material was constituted of small satellite colonies which might serve as a source for growth reinitiation. Accordingly, the growth rate reflected by dry weight increase upon glucose resupply was higher in *wt* cultures than in *frq*^10^ and *Δwc-1* (Figure 5B, After, 5C). Filamentous fungi naturally grow on solid surfaces. Hence, growth regeneration was tested also on solid media by comparing growth rates following the transfer of mycelia from starved and control liquid cultures onto race tube medium which contained glucose (Figure 5D, left panel). While the growth rate of starved *wt* cultures approached that of the non-starved controls within 28 hours, it remained at a relative low level in *Δwc-1* (Figure 5D, right panel). Because of its low growth rate on solid medium *frq*^10^ could not be tested under these conditions.

**Figure 5.**
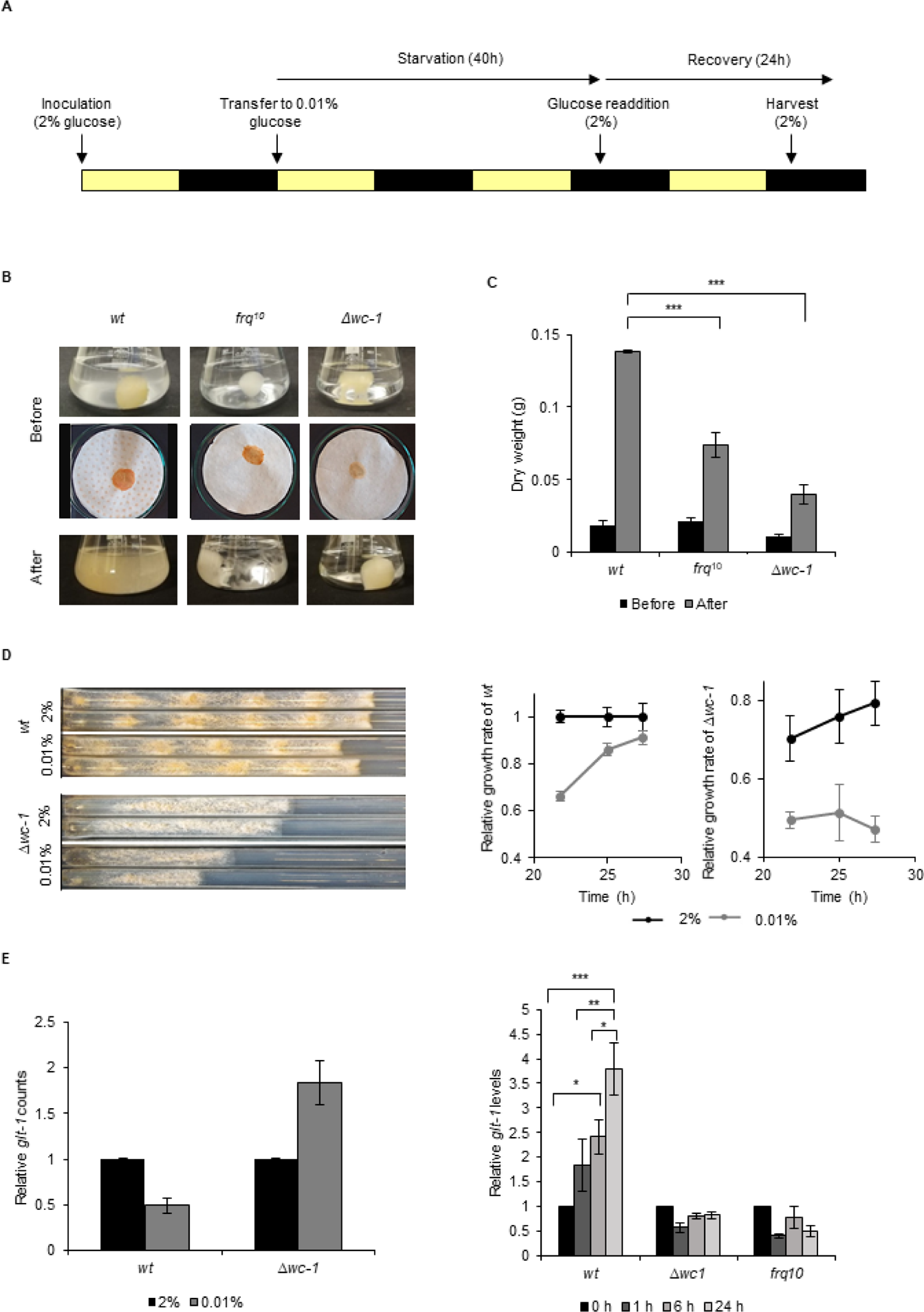
Proper recovery from starvation requires a functional clock. (A) Schematic design of the experiment. Mycelial discs of the *wt* and the clock mutant strains were incubated in standard liquid medium in L/D12. After 24 hours mycelial balls were transferred to starvation medium. Following 40 hours of starvation, glucose was added to the medium. Yellow and black bars indicate the periods cultures spent in light and darkness, respectively. (B) Comparison of the growth of *wt* and clock mutants after glucose resupply. Pictures of the liquid (before: upper images) and the vacuum filtered cultures (before: lower images) were taken after 40 hours of starvation and 24 hours after glucose resupply (after). (C) *wt* grows faster after glucose supplementation than clock mutants. Dry weight of cultures was measured at the indicated phases of the experiment. (n=3, ±SEM, Factorial ANOVA, significant strain*treatment interaction, post hoc analysis: Tukey HSD test) (D) Growth of *wt* responds effectively to glucose supplementation also on solid media. Experiments were performed as described in A. However, following the 40 hours incubation in liquid medium, mycelia were transferred to solid race tube medium containing 0.2% glucose. Race tube images (left panel) and quantification of their growth rate (right panels) are shown. (n=4-6) 0.01% and 2% indicate the glucose content in the liquid medium. Growth rates were normalized to that of *wt* samples grown in standard (2% glucose) liquid medium before the transfer onto solid medium. (E) Lack of the functional clock affects the proper alignment of glucose transporter expression to glucose levels. Left panel: *glt-1* counts in RNAseq data. Values were normalized to that of cultures grown in 2% glucose (time point 0: start of starvation). Right panel: Experiments were performed as described in (A). Samples were harvested at the indicated time points following glucose readdition and relative levels of *glt-1* RNA were determined by qPCR. Values were normalized to that of time point 0 (end of starvation, start of recovery) (n=3, ±SEM, Factorial ANOVA, significant strain*treatment interaction, post hoc analysis: Tukey HSD test)

In *Neurospora* a dual-affinity transport system mechanism enables the proper adaptation of glucose uptake to external glucose levels. When glucose levels are elevated, increased expression of the low-affinity and high-capacity transporter, *glt-1* ensures efficient utilization of the carbon source (Wang et al., 2017). Accordingly, the transcriptomic analysis revealed higher *glt-1* levels in non-starved than starved *wt* cells. In *Δwc-1*, however, *glt-1* was upregulated under starvation conditions, indicating that the WCC is crucial in the adaptation of glucose transport to nutrient levels (Figure 5E, left panel). Next, we examined *glt-1* expression during recovery. In *wt*, *glt-1* RNA levels gradually increased upon supplementation of the starvation medium with 2 % glucose, whereas no significant expression changes were detected in either *Δwc-1* or *frq*^10^ (Figure 5E, right panel). In summary, the marked differences between the recovery behaviour of the different strains suggest that adaptation to changing nutrient availability is more efficient when a circadian clock operates in the cell.

## Discussion

In nature, the reproductive capacity of living things exceeds the availability of resources, and organisms face nutrient shortages from time to time. Optimal metabolic responses to nutrient deficiencies could therefore provide a significant advantage in selection. Deprivation of carbon sources significantly reduces the transcription/translation rate, which is also a challenge for the TTFL-based molecular clock. Nutrient compensation of the circadian rhythm and thus timely regulation of physiology could therefore be a prerequisite for appropriate adaptation. Short glucose deprivation does not affect expression of the core clock proteins and this maintenance of the protein levels was considered essential for nutrient-compensation (Emerson et al., 2015; Gyongyosi et al., 2017; Sancar et al., 2012). Our experiments show that the circadian TTFL can function with substantially altered levels and stoichiometry of its key elements. Thus, WCC levels decrease sharply during glucose starvation and FRQ becomes hyperphosphorylated, while the clock maintains the period of expression of *frq* and various output genes. Cytosolic but not nuclear WC protein levels are primarily reduced under low glucose conditions. Cytosolic WCC might serve as a reserve pool of the positive factor that can be rapidly activated by Zeitgebers (e.g. light) to synchronize the clock with the changing environment (Malzahn et al., 2010). In accordance with this, we found that acute light response is reduced under starvation.

As RNA levels encoding the WCC subunits remain compensated during glucose starvation, posttranscriptional mechanisms may be involved in the reduction of their protein levels. Hyperphosphorylation of FRQ can weaken the feedback on the WCC, which might contribute to the compensation for the reduced transcription/translation rate during nutrient shortage and can lead to faster degradation of the WCC (Figure 6). The starvation-dependent reduced stability of WC-1 in *wt* supports this notion. The absence of FRQ-dependent feedback on WCC might explain why WCC levels are only moderately affected by starvation in *frq*^9^. The starvation-dependent decrease in *frq*^9^ RNA abundance also suggests that in the absence of the feedback mechanism, the starvation-induced decrease in transcription is not compensated by an increase in WCC activity. The glucose-dependent modulation of the clock is probably not restricted to changes in the FRQ-dependent recruitment of CK1a to the WCC, as the impaired interaction between FRQ and CK1a in the *frqΔFCD1-2* strain only partially affected WCC level alterations and did not prevent the starvation-induced hyperphosphorylation of FRQ.

**Figure 6.**
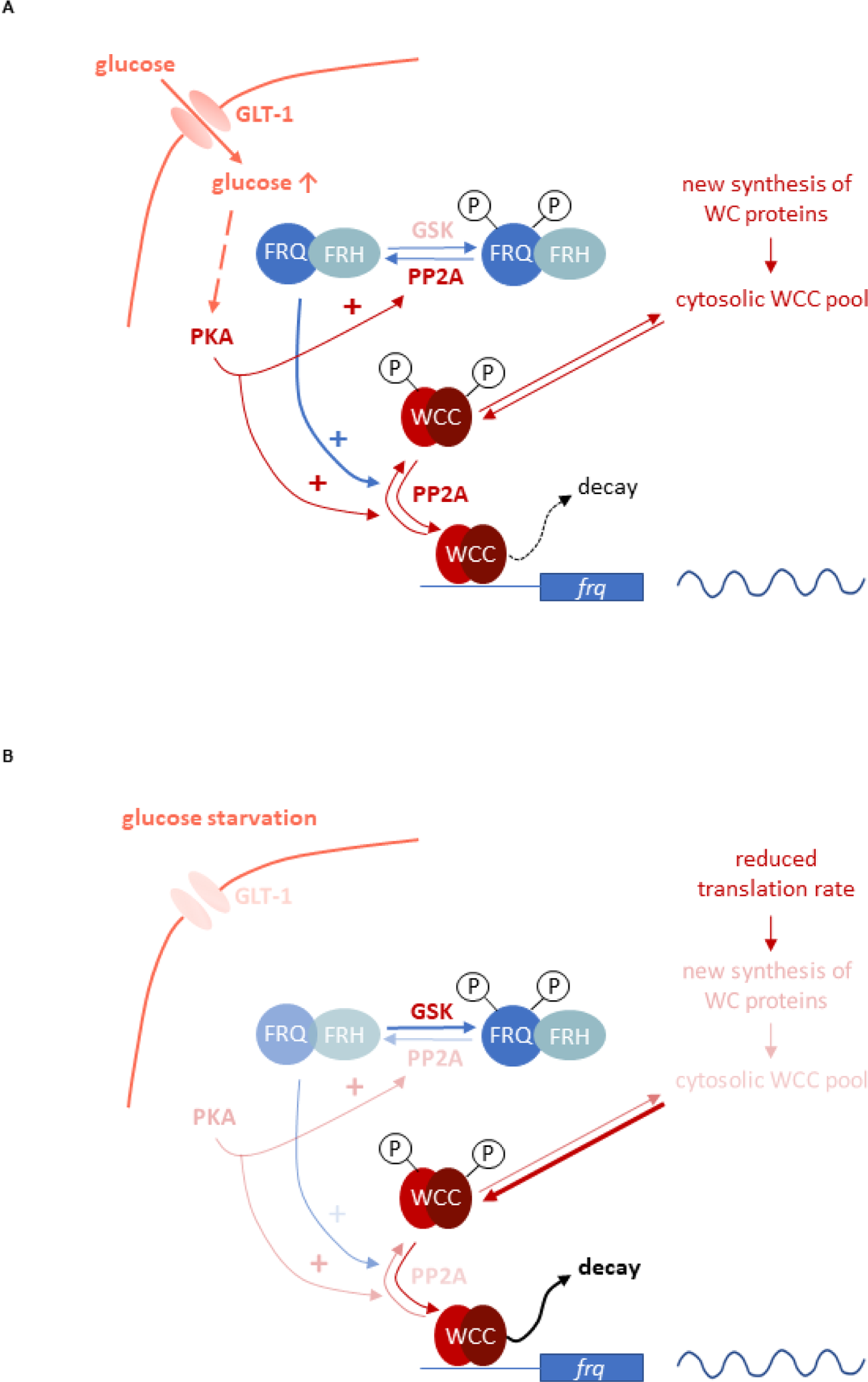
Model representing the role of the negative feedback and the PKA-, PP2A- and GSK-mediated signaling in the control of the molecular clock at high glucose levels (A) and under starvation (B). Starvation reduces the activity of both PKA and PP2A but stimulates GSK. PKA can act as a central regulator of the starvation-induced modifications of the clock components, as its weakened activity results in enhanced action and the consequent destabilization of the WCC, resulting in compensated *frq* transcription at significantly reduced WC levels. PKA also affects PP2A. Reduced activity of PP2A and the parallel induction of GSK in starvation can lead to hyperphosphorylation of FRQ which in turn lessens the negative feedback on the WCC. Higher and lower activities of enzymes and processes are indicated by more and less intense colours, respectively.

Starvation triggers characteristic changes in the activity of signaling pathways that affect basic components of the circadian clock. We found that PP2A, GSK and PKA are involved in the starvation-induced modification of the molecular clock (Figure 6). PP2A was described as glucose sensitive regulator in fungal as well as in mammalian cells (Hallett et al., 2014; Lee et al., 2018) and we showed that glucose starvation decreases PP2A activity in *Neurospora* as well. PP2A is known to act on both the negative and the positive element of the *Neurospora* clock (Schafmeier et al., 2005; Yang et al., 2004). Reduced PP2A activity during nutrient limitation may contribute to the hyperphosphorylation of FRQ lessening the negative feedback on the WCC activity. However, this effect might be antagonized by a weaker dephosphorylation of the WC proteins. In the *rgb-1^RIP^* strain in which one of the regulatory subunits of PP2A is disrupted and the enzyme activity is rather low, WC levels did not respond to glucose-deprivation, suggesting that starvation-dependent expression change of the WCC is dependent on PP2A. Starvation triggers activation of GSK in many organisms. Our data indicate that hyperphosphorylation of FRQ upon glucose depletion is dependent on GSK. Whether FRQ is a direct substrate of GSK, is still to be elucidated. Reduced PKA activity in starvation is considered as a central organizer of the cellular adaptation to nutrient restriction (Huang et al., 2007; Li and Borkovich, 2006a). PKA is a priming kinase for subsequent phosphorylation of the WCC by CK1a and/or CK2, contributing to inactivation of the transcription factor. However, since PKA increases FRQ stability (Huang et al., 2007), affects translation under stress conditions (Barraza et al., 2017; Leipheimer et al., 2019) and activates PP2A in yeast (Castermans et al., 2012), it may also affect the clock via these functions. Indeed, the enhanced PKA activity in the *mcb* strain interfered with all observed starvation-induced changes within the molecular oscillator. In conclusion, our observations along with literature data suggest a model in which PKA acts as a main coordinator in the adaptation of the circadian clock to different nutrient conditions (Figure 6). Overall, the nutrient-dependent activity of multiple signaling pathways might ensure that the transcriptional activity of the WCC and thus expression and oscillation of *frq* remain nutrient-compensated. Indeed, long-term starvation had an effect on more than 2000 RNAs, indicating global changes in cell functions that might affect diverse regulatory systems.

Presence of a functional circadian clock significantly affects the adaptation of *Neurospora crassa* to long-term glucose deprivation at both the phenotypic and transcriptomic level. We showed that starved *wt* cultures better adapt the expression of the high-capacity glucose transporter to glucose levels and grow faster upon resupply of glucose than clock-less cultures. Importantly, more than 1300 coding genes displayed glucose specific differences in the transcriptome response between *wt* and *Δwc-1*. These genes affect various cell functions including carbohydrate, amino acid and fatty acid metabolism. The lack of a functional WCC resulted in a characteristic shift from the control of central carbon metabolism and the downregulation of amino acid synthesis to the upregulation of amino acid catabolism and the downregulation of fatty acid synthesis. In conclusion, our results suggest that the WCC has a major impact on the mechanisms balancing the cellular energy state and highlights the importance of the circadian clock in cell survival under nutritional stress and strengthens the evolutionary significance of circadian timekeeping.

## Materials and Methods

### RESOURCE AVAILABILITY

#### Lead contact

Further information and requests for resources and reagents should be directed to and will be fulfilled by the lead contact, Krisztina Káldi.

#### Materials availability

Plasmids and strains generated in this study are available upon request from the lead contact.

#### Data and code availability

Data generated in this study are available from the corresponding author on request.

### EXPERIMENTAL MODEL AND SUBJECT DETAILS

*Neurospora crassa* strains *wt* (FGSC #2489), *wt,bd* (FGSC #1858), *frq^10^,bd* (FGSC #7490), *frq^9^,bd* (FGSC #7779), *rgb-1^RIP^* (FGSC #8380), *mcb* (FGSC #7094) and *Δprd-4* (FGSC #11169) were obtained from the Fungal Genetics Stock Center (http://www.fgsc.net/ (McCluskey, 2003)). The last two knockouts were created during the Neurospora Genome Project (Colot et al., 2006). *Δwc1,bd, frq^10^,bd,his-3* (Bell-Pedersen et al., 1996) and *qa-gsk* (Tataroglu et al., 2012) strains were generous gifts from M. Brunner. To generate the *frq^10^,Δfcd1-2* strain, *pBM60-ClaI-ΔFCD1-2* (Querfurth et al., 2011) (generous gift from M. Brunner) was integrated into the *his-3* locus of *frq^10^ bd,his-3* by electroporation (Margolin, 1997).

The standard liquid medium contained Vogel’s medium (Vogel, 1964) supplemented with 0.5% L-arginine, 10ng/ml biotin and 2% of glucose. In the starvation medium the glucose concentration was reduced to 0.01%. For liquid cultures mycelial mats were grown in Petri dishes in standard liquid medium for two days at room temperature in constant darkness. From the mycelial mat discs were punched out, washed by sterile distilled water and transferred into Erlenmeyer flasks containing liquid medium and were grown at 25°C and shaken with 90rpm if not indicated otherwise. For growing, at least 150 ml of liquid medium per mycelial discs was used in order to keep glucose concentration as constant as possible during the incubation. Under these conditions *Neurospora* grows as balls of mycelium.

## METHOD DETAILS

### Protein Analysis

Extraction of *Neurospora* protein, Western blots (Gyongyosi et al., 2013; Schafmeier et al., 2006) and subcellular fractionation (Gyongyosi et al., 2017; Luo et al., 1998) was performed as described earlier. In subcellular fractionation same amount of cytosolic and nuclear fractions were analyzed. PonceauS staining was used as loading control. Representative Western-blots are shown. Protein levels were determined by densitometry with the ImageJ software (https://imagej.nih.gov/ij/download.html).

### RNA Analysis

Total RNA was extracted using the TriReagent^®^ (Sigma Aldrich #93289) isolation reagent and transcript levels were quantified as described earlier (Gyongyosi et al., 2013). Values were normalized to the *wt* grown in 2% glucose containing standard medium in each experiment unless indicated otherwise. As level of previously used housekeeping gene *actin* decreased under starvation conditions, we chose new reference genes for our experiments based on previous data (Cusick et al., 2014; Hurley et al., 2015; Llanos et al., 2015). We found *gna-3* to be the least variable under the applied experimental conditions. Ratios of reference genes are presented in Figure S10. For sequences of oligonucleotides and hydrolysis probes see Key resources table.

### RNA Sequencing and Data Analysis

Liquid cultures were grown under standard and glucose starving conditions in 12-12 hours light-dark cycles for 48 hours and samples were harvested at ZT12 (n=4 for each group). RNA was purified using the TriReagent^®^ (Sigma Aldrich #93289) isolation reagent. Following DNase treatment RNA quality was controlled by the Nanodrop^TM^ One^C^ spectrophotometer, the Qubit^TM^ 4.0 Fluorometer (Invitrogen) and the Agilent TapeStation 4150 System.

Library preparation and sequencing were performed by BGI Genomics, China, using PE-100 library. Sequencing quality check was performed with FastQC (Andrews, 2010). Mapping was performed with STAR (Dobin et al., 2013) to the *N.c*. genome from Ensembl (Neurospora_crassa.NC12.48) (Yates et al., 2020), and the indexing and duplicate filtering with samtools (Li et al., 2009). Read counting was done using HTSeq-count (Anders et al., 2015). Differential expression analysis was done with DESeq2 (Love et al., 2014) package in R (R Core Team, 2020). (Raw data of the RNA sequencing: https://doi.org/10.5061/dryad.t4b8gtj4p) For primer sequences used during the validation of RNAseq results see Key resources table. Regarding Figures S6-S8 and Tables S3 it is important to note that KEGG analysis annotates less genes to the different metabolic pathways than GO analysis.

### Phosphatase Assay

Measuring PP2A activity was carried out with a Ser/Thr Phosphatase Assay System (Promega #V2460) according to the manufacturer’s instructions. Each reaction contained 10 μg protein and was performed for 20 minutes at room temperature. Absorbance was measured at 600nm.

## QUANTIFICATION AND STATISTICAL ANALYSIS

For statistical analysis, the Statistica 13 (Statsoft Inc., Tulsa, OK, USA) software was used. Error bars indicate ±SEM. Results were considered to be statistically significant when p value was <0.05. (*p<0.05; **p<0.01; ***p<0.001; n.s.: non-significant) Further statistical details can be found in the figure legends. N represents the number of independent biological samples.

## Acknowledgements

We thank the Fungal Genetics Stock Center and the Neurospora Genome Project for providing *Neurospora* strains. We also thank Rita Krisztina Nagy and Fanni Kóródi for excellent technical assistance and Ágnes Réka Sűdy for critical reading of the manuscript. This work was supported by grants of the National Research, Development and Innovation Office, NKFIH (K115953 and K132393 to K.K.; FK132474 to N. G. and FK134267 to O.K) and by the ÚNKP-19-3-IV-SE-3 New National Excellence Program of the Ministry for Innovation and Technology to A. Sz..

## Author contributions

Conceptualization, K. K., N. G., O. K. and M. B.; Investigation and Analysis A. S., N. G., O. S. and A. R. C. D.; Resources and Funding Acquisition, K. K. and N. G.; Data Curation, G. S., K. K., N. G. and A. S.; Writing and editing, K. K., N. G., A. S., M. B.

## Competing interests

The authors declare no competing interests.

---

**Supplemental Figure 1.**
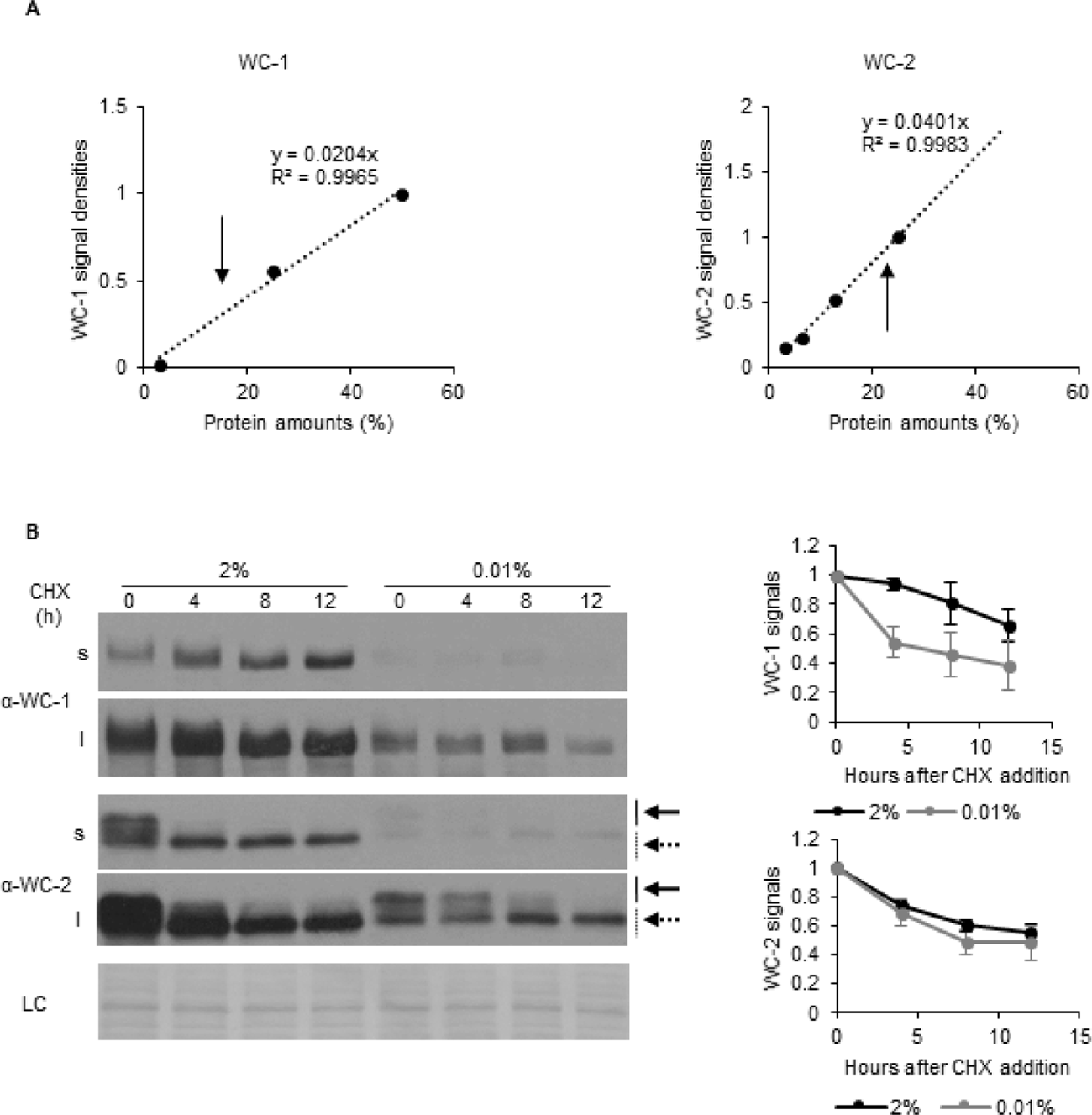
Analysis of expression changes of clock components. (A) Calibration of the WCC expression levels in starvation. Protein gel was loaded with increasing quantities (3,13%-50%) of wt lysates grown in 2% glucose containing medium and analyzed by Western blotting. WC-1 (left panel) and WC-2 (right panel) specific signals were analyzed by densitometry and a calibration line was fitted by linear regression. Arrows indicate the amount of positive clock components in wt lysates grown in starvation medium. (Figure 1B) (B) Glucose starvation moderately reduces the stability of WC-1. Experimental procedures were performed as described in Figure 1C. Following 40 hours of starvation translation inhibitor cycloheximide (CHX) was added to the medium in a final concentration of 10µg/ml (time point 0). Samples were harvested at the indicated time points following CHX addition. Left panel: Whole cell extracts were analyzed by Western blotting with the indicated antibodies. Solid and dashed arrows indicate hyper- and hypophosphorylated forms of WC2 proteins, respectively. Short (s) and long (l) exposures are shown in order to get comparable signals at both conditions. (LC: loading control) Right panel: Signal densities of WC proteins were determined and normalized to the values detected in time point 0. (n=5, ±SEM).

**Supplemental Figure 2.**
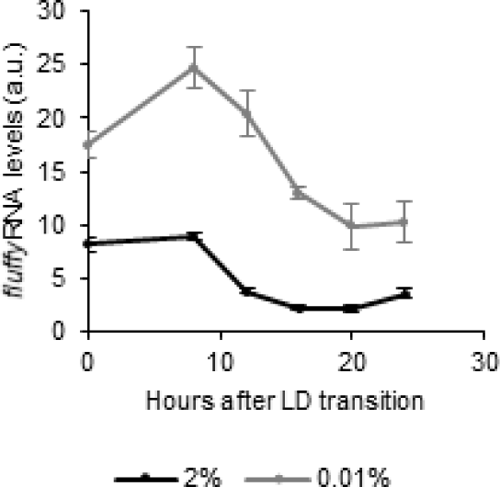
The circadian output is rhythmic in starvation. Following an incubation in standard liquid medium for 24 hours, mycelia were transferred to fresh standard or starvation medium. After 24 hours of incubation in LL, cultures were transferred to DD. Samples were harvested at the indicated time points. Relative *fluffy* RNA levels were determined. (n=3-6, ±SEM)

**Supplemental Figure 3.**
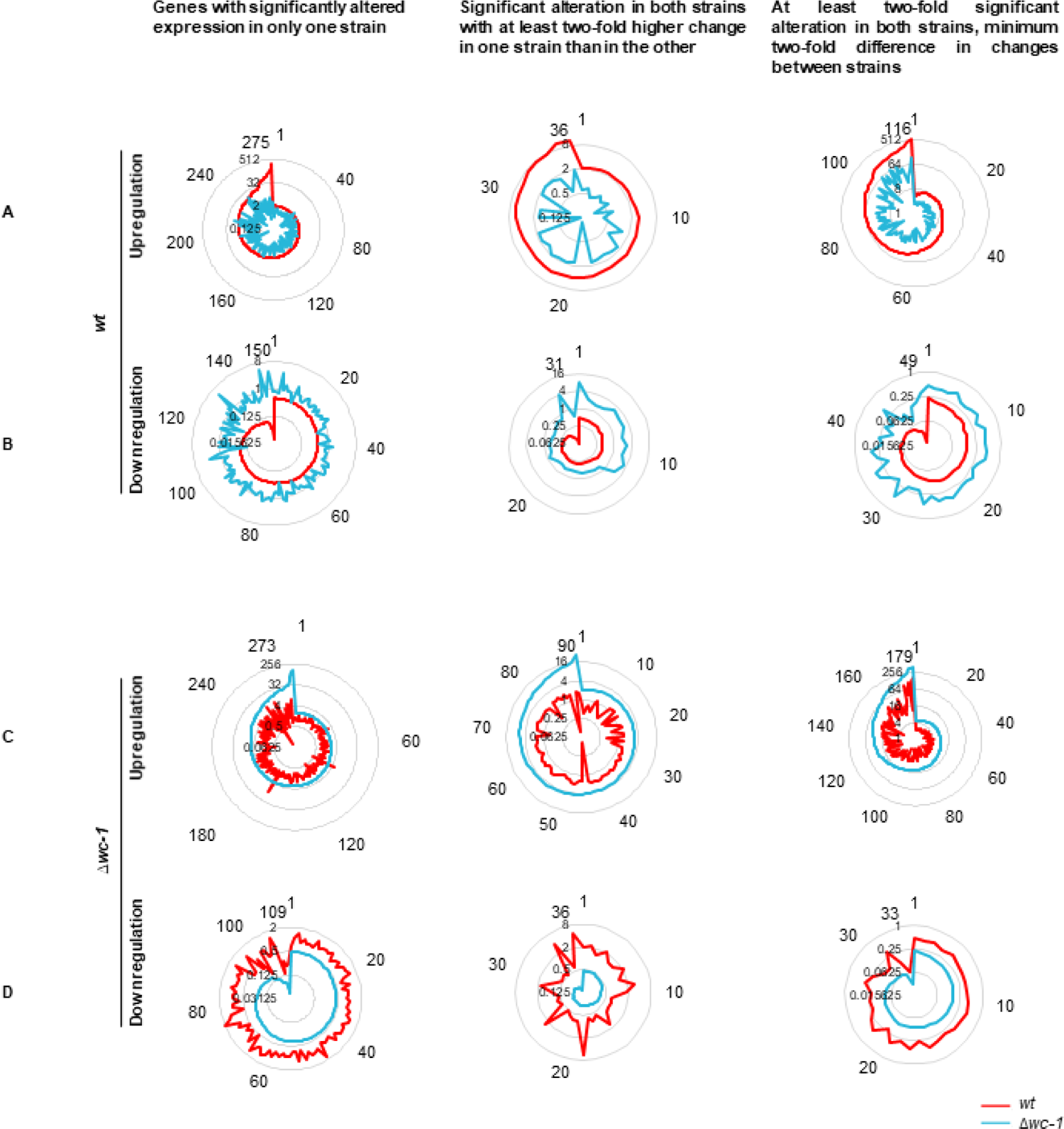
Strain-specific differences of gene expression changes in response to starvation. Concentric circles indicate the fold-change, numbers around the diagram indicate the genes ordered according to the extent of up/downregulation. A: Genes showing *wt*-specific increase in their RNA levels in response to starvation. B: Genes showing *wt*-specific decrease in their RNA levels in response to starvation. C: Genes showing *Δwc-1*-specific increase in their RNA levels in response to starvation. D: Genes showing *Δwc-1*-specific decrease in their RNA levels in response to starvation.

**Supplemental Figure 4.**
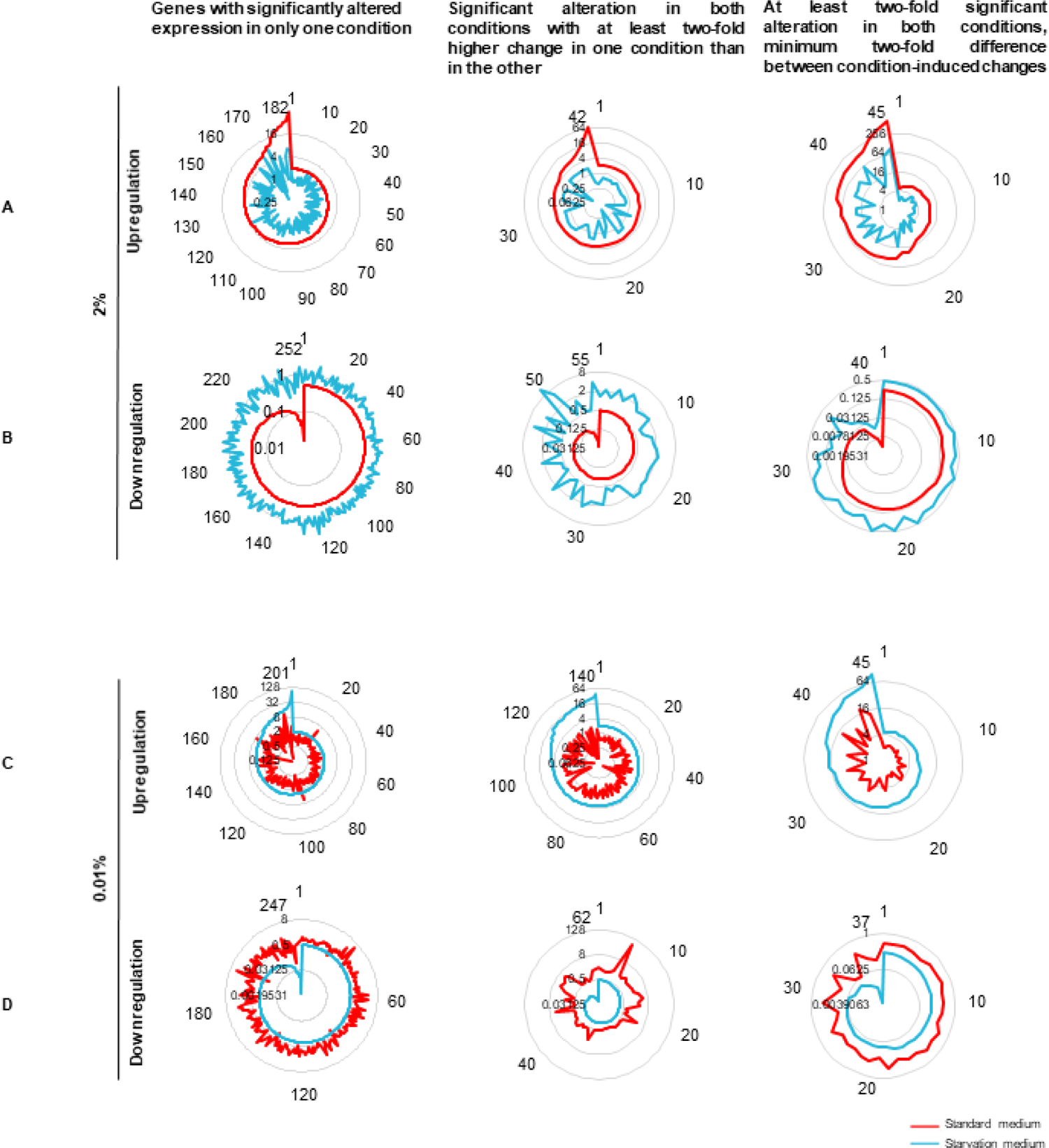
Glucose-specific differences in Δwc-1/wt ratio of gene expression. Concentric circles indicate the fold-change, numbers around the diagram indicate the genes ordered according to the extent of up/downregulation. A: Genes showing glucose-specific (2%) increase in their *Δwc-1/wt* ratio. B: Genes showing glucose-specific decrease in their *Δwc-1/wt* ratio. C: Genes showing starvation-specific (0.01%) increase in their *Δwc-1/wt* ratio. D: Genes showing starvation-specific (0.01%) decrease in their *Δwc-1/wt* ratio.

**Supplemental Figure 5.**
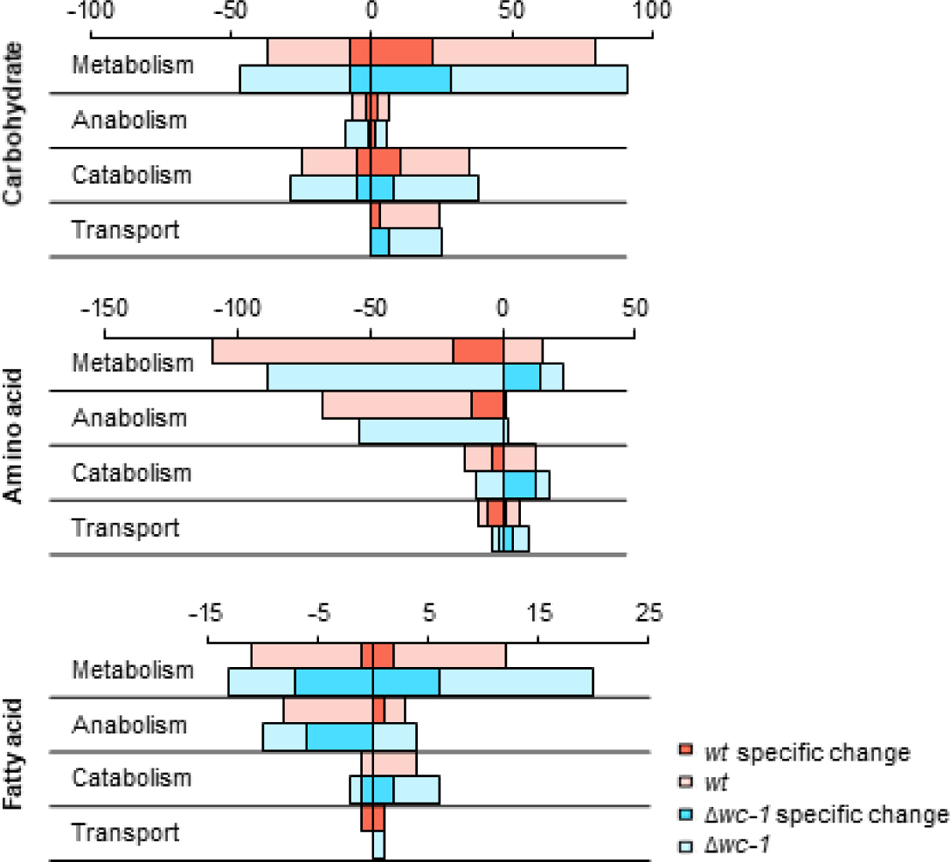
Characterization of different metabolic pathways in strain-specific responses to glucose starvation. Expression changes of genes involved in amino acid, carbohydrate and fatty acid metabolism in response to 40-hour glucose starvation. Positive values show number of genes with increased, while negative values indicate number of genes with decreased RNA levels. Number of genes showing strain-specific changes in *wt* or *Δwc-1* are marked with darker red and blue color, respectively. Genes were classified by GO analysis (Mi et al., 2013).

**Supplemental Figure 6.**
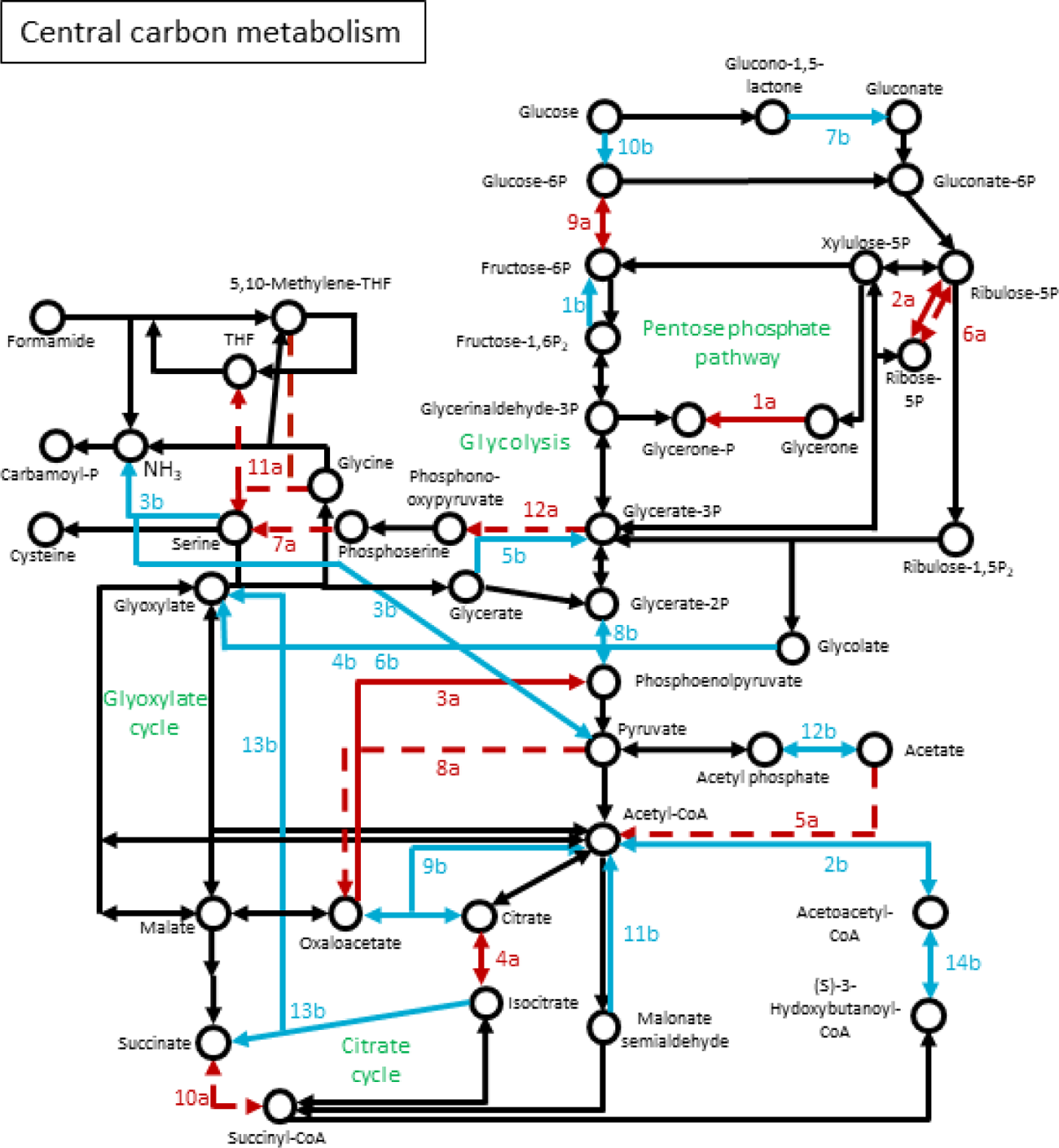
Genes of central carbon metabolism, that showed strain-specific change to starvation. Network of genes were constructed based on the KEGG Mapper tool. Numbering of genes is resolved in Supplemental table 3. Red arrow: *wt*-specific change; Blue arrow: *Δwc-1* specific change; Solid line: significant increase; Dashed line: significant decrease

**Supplemental Figure 7.**
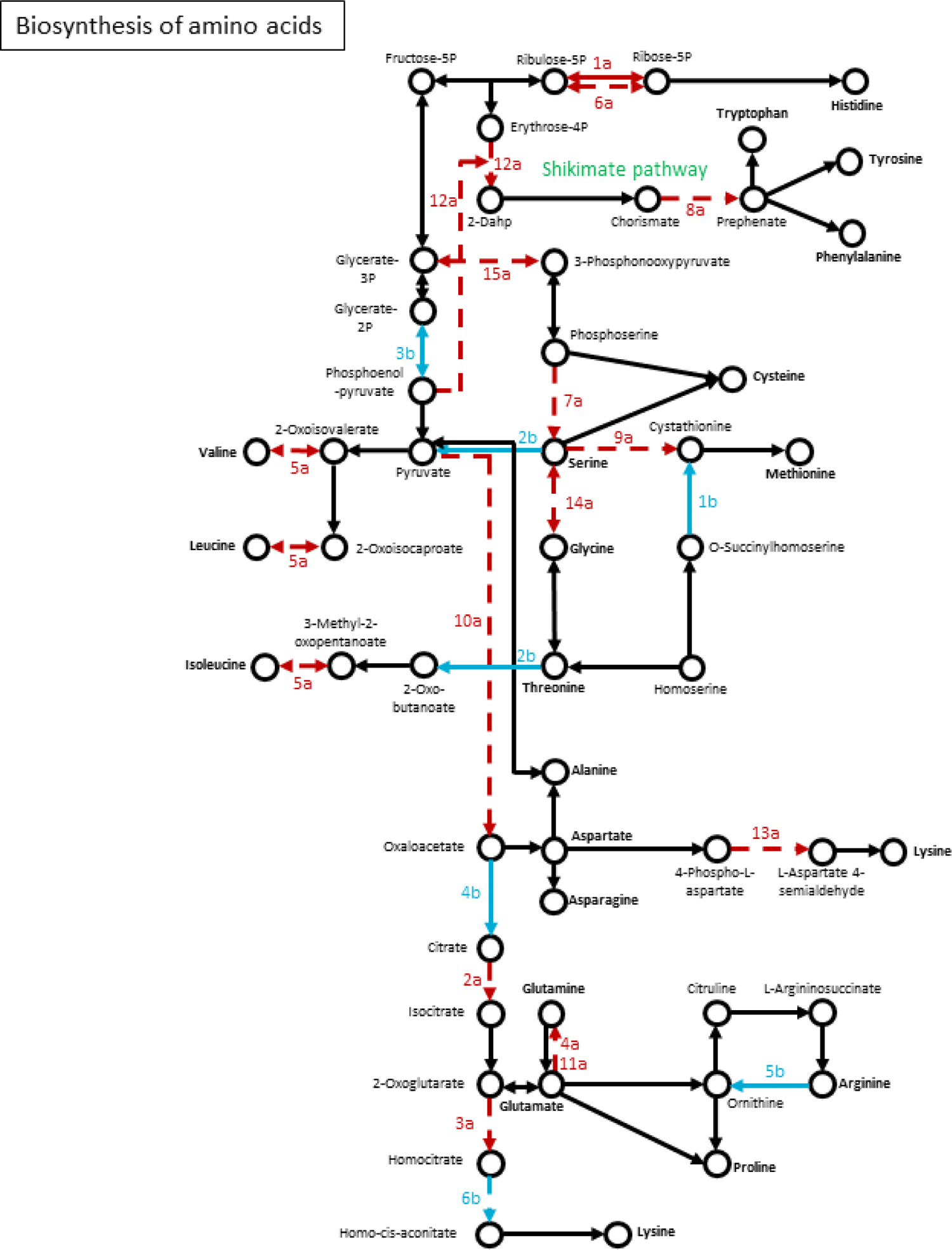
Genes of amino acid biosynthesis, that showed strain-specific change to starvation. Network of genes were constructed based on the KEGG Mapper tool. Numbering of genes is resolved in Supplemental table 3. Red arrow: *wt*-specific change; Blue arrow: *Δwc-1* specific change; Solid line: significant increase; Dashed line: significant decrease

**Supplemental Figure 8.**
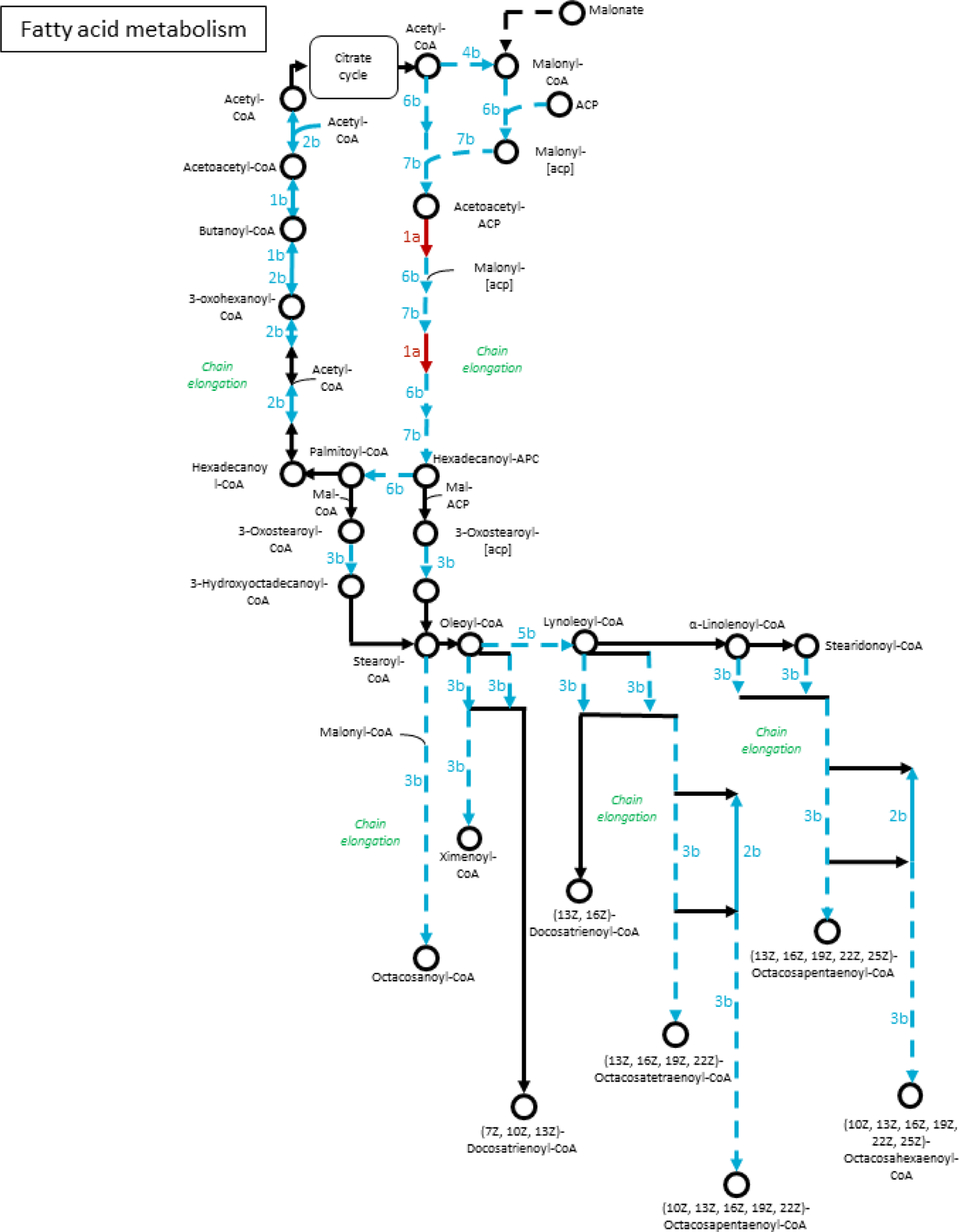
Genes of fatty acid metabolism, that showed strain-specific change to starvation. Network of genes were constructed based on the KEGG Mapper tool. Numbering of genes is resolved in Supplemental table 3. Red arrow: *wt*-specific change; Blue arrow: *Δwc-1* specific change; Solid line: significant increase; Dashed line: significant decrease

**Supplemental Figure 9.**
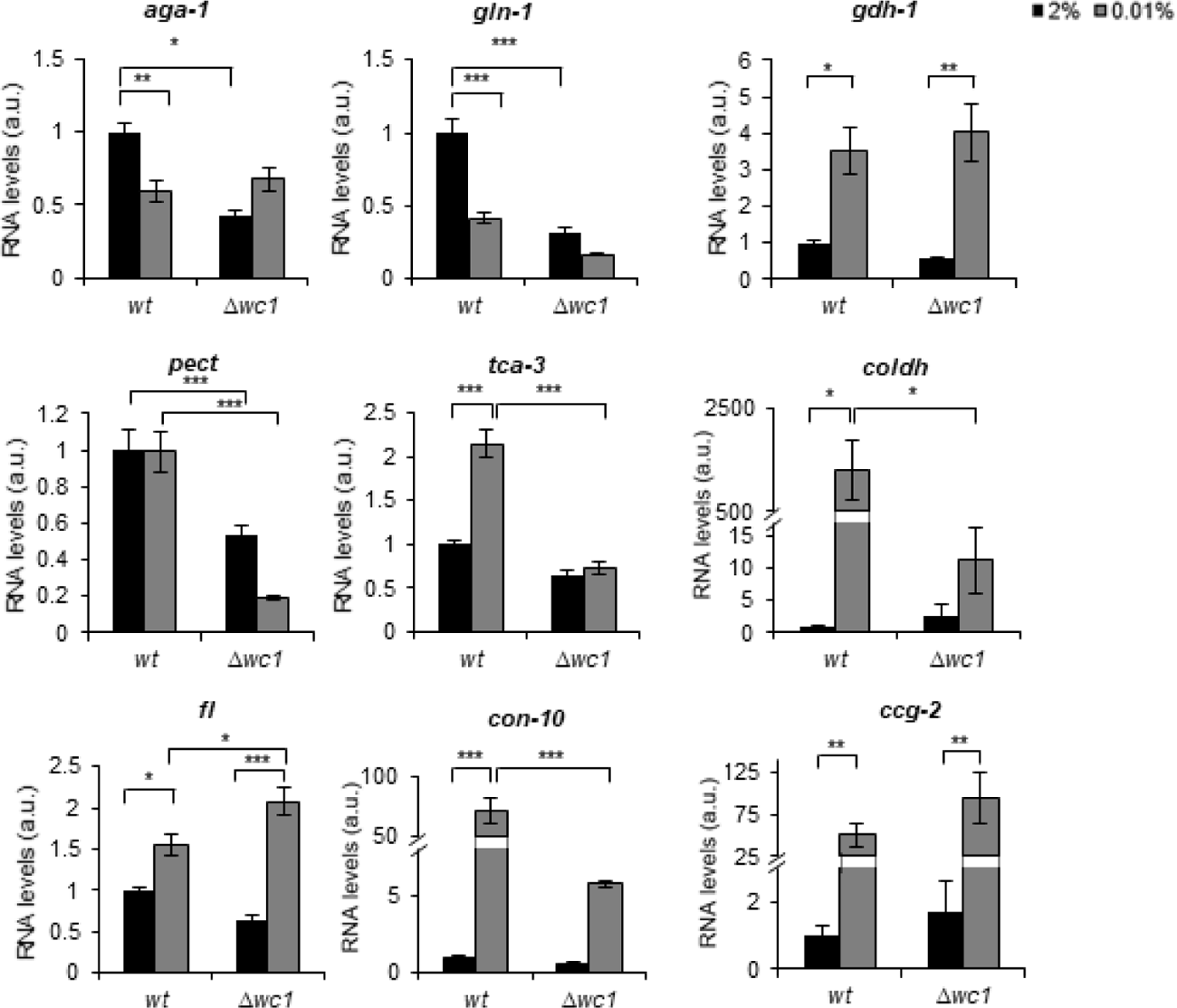
qPCR validation of the RNA-seq data. Three groups of processes (amino acid metabolism, carbohydrate metabolism and conidiation) that were hypothesized to change in response to glucose deprivation were tested. Values were normalized to those measured in *wt* grown in standard medium. (*aga-1*: arginase (NCU02333); *gln-1*: glutamine synthetase (NCU06724); *gdh-1*: NAD-specific glutamate dehydrogenase (NCU00461); *pect*: pectin esterase (NCU10045); *tca-3*: aconitate hydratase, mitochondrial (NCU02366); *choldh*: choline dehydrogenase (NCU01853); *fl*: conidial development protein fluffy (NCU08726); *con-10*: conidiation-specific protein 10 (NCU07325); *ccg-2*: hydrophobin (NCU08457)) (n=4, ±SEM, Factorial ANOVA, significant treatment effect (*gdh-1, ccg2*), significant strain effect (*pect*) significant strain*treatment interaction (*aga-1, gln-1, tca-3, coldh, fl, con-10*))

**Supplemental Figure 10.**
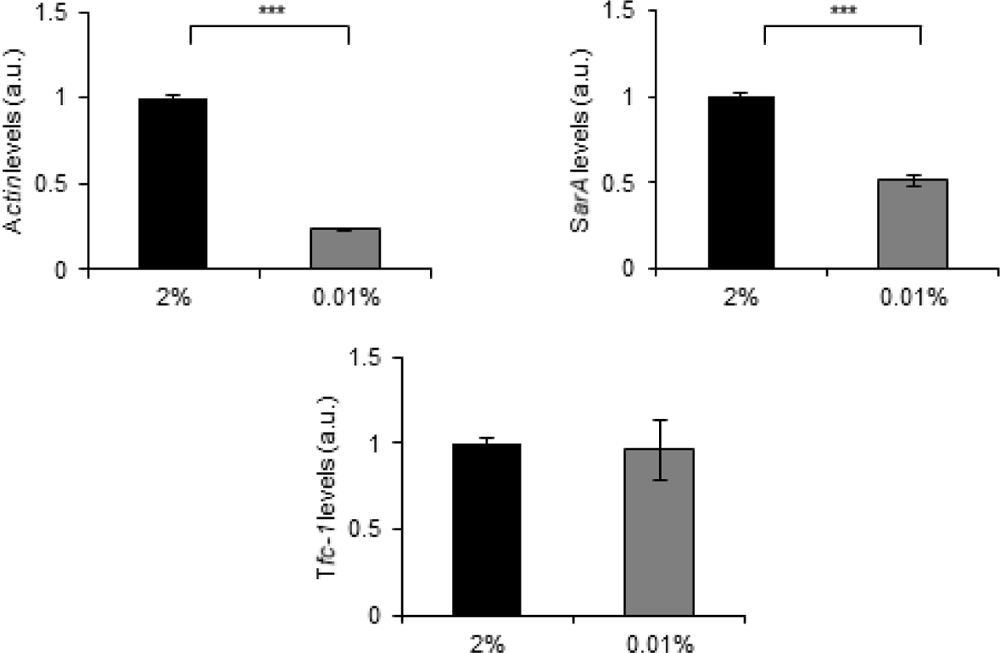
Actin levels are decreased in glucose starvation. Experimental procedures were performed as described in Figure 1C. Transcript levels of the indicated genes were quantified by qPCR relative to *gna-3* levels. (n=4; two-sample t-test;)

**Supplemental Table 2.**
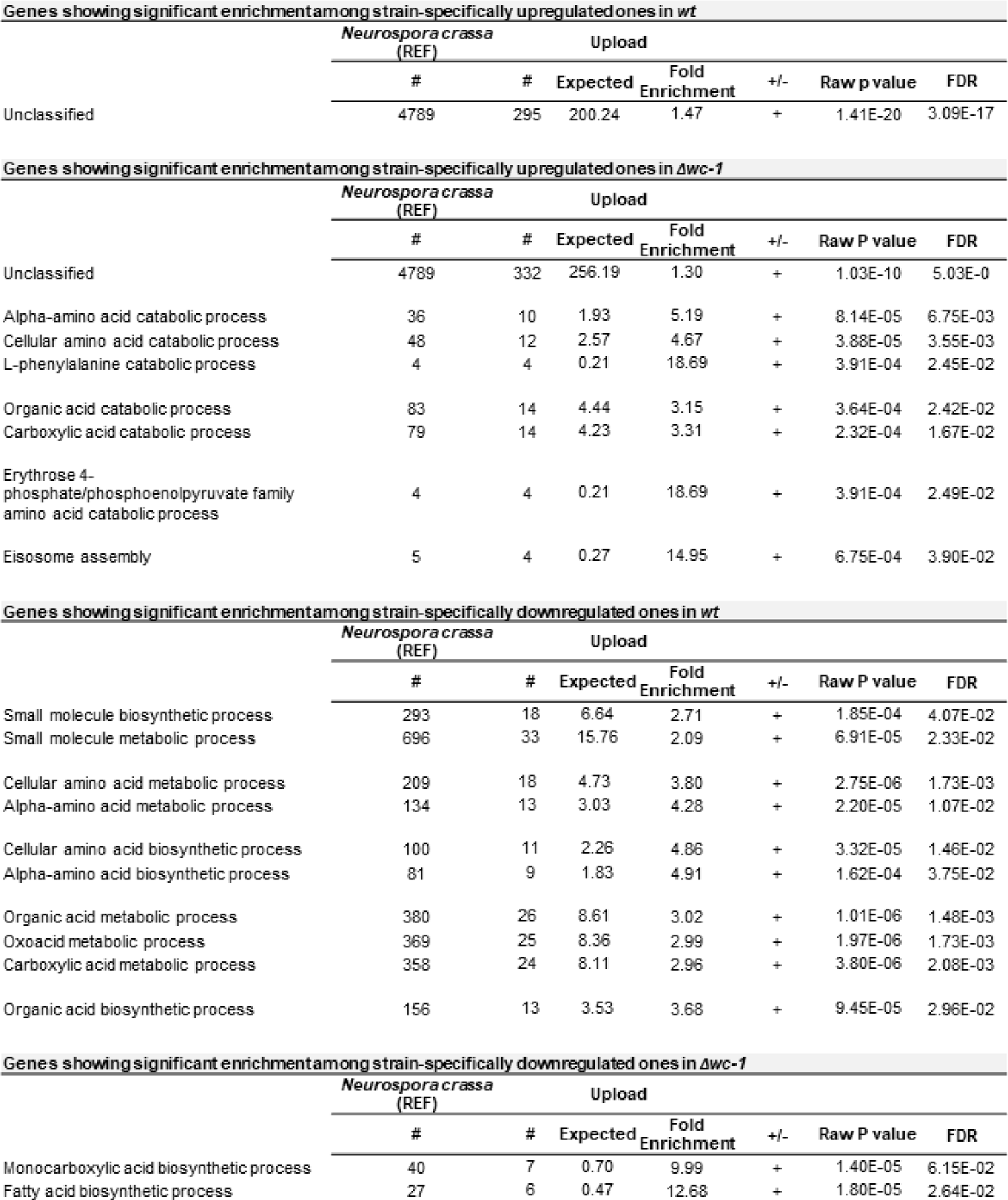
*Gene Ontology (GO) enrichment analysis of genes showing strain-specific response to starvation*. Significantly enriched functions are shown. Go enrichment analysis was performed on data obtained by the analysis shown in Supplemental Figure 3, i.e. on genes showing exclusive or significantly higher change in their expression rate in one of the strains. (FDR: false discovery rate)

**Supplemental Table 3.**
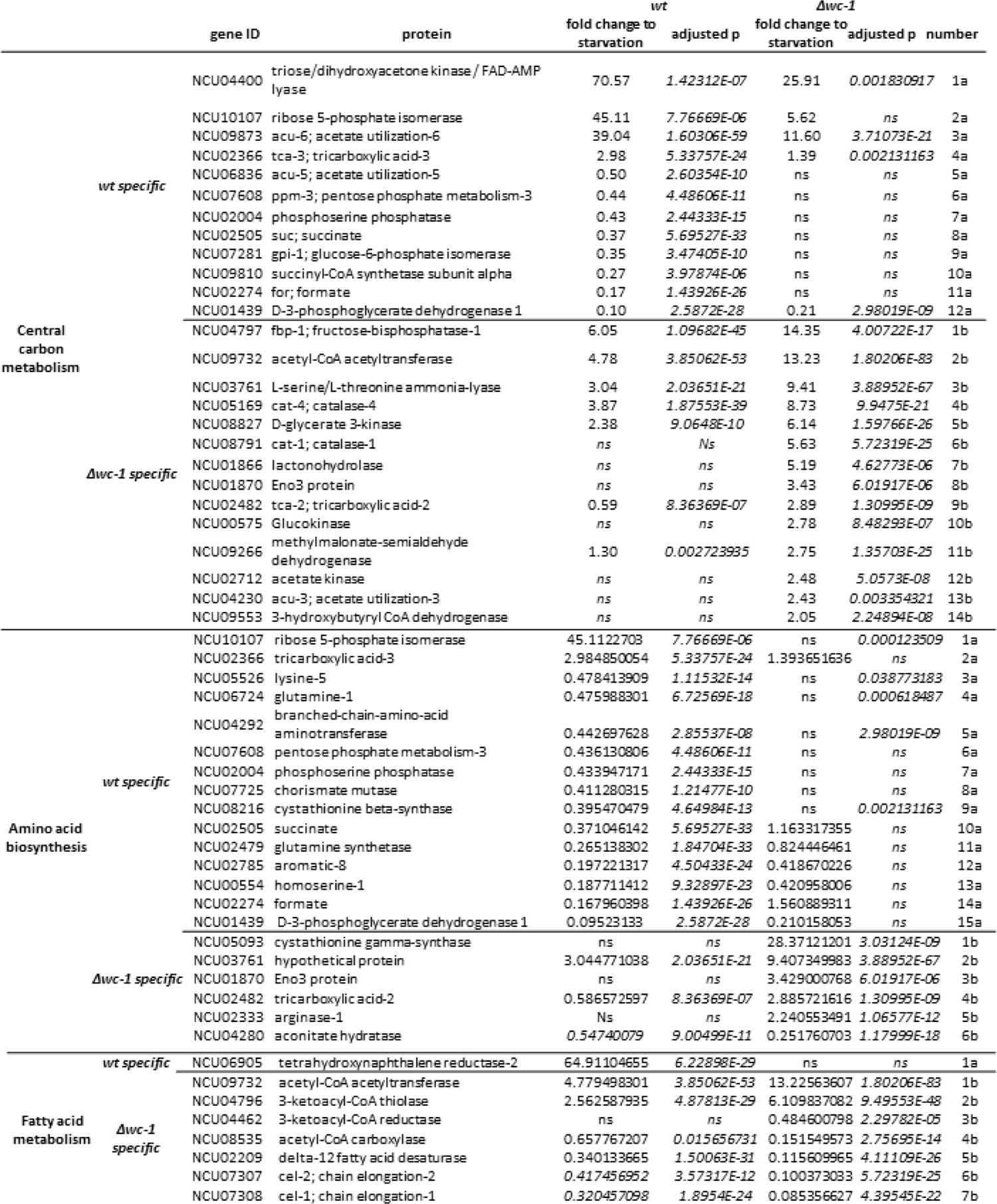
*Genes of central carbon metabolism amino acid biosynthesis and fatty acid metabolism, that showed strain-specific expression change to starvation*. Genes were selected with the help of the KEGG Mapper tool. Numbering of genes in Supplemental Figure 6-8 can be found in the last column.

**Supplemental Table 4.**
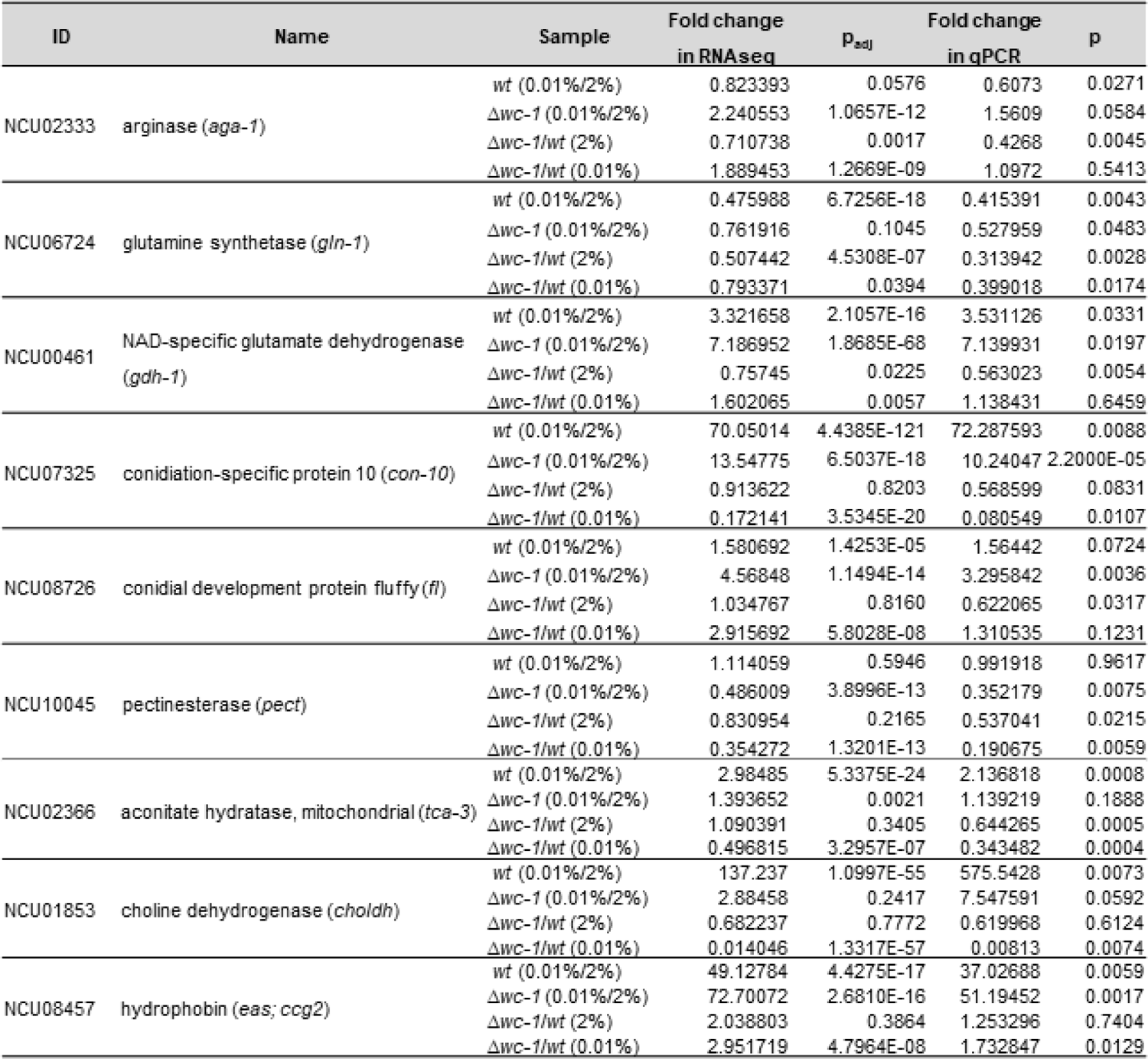
Comparison of the results from RNA-seq and the experimental validation of the chosen genes with qPCR. Experimental procedures were performed as described in Figure S9. (n=4, two sample t-test)

## SOURCE DATA LEGENDS

### Source data for Figure 1 B

**Figure 1 B – source data FRQ:** Western blots were used to detect expression of FRQ in the indicated samples for Figure 1 B.

**Figure 1 B – source data FRQ_labeled:** Figure with the area highlighted was used to develop the Figure 1 B for FRQ.

**Figure 1 B – source data WC1:** Western blots were used to detect expression of WC1 in the indicated samples for Figure 1 B.

**Figure 1 B – source data WC1_labeled:** Figure with the area highlighted was used to develop the Figure 1 B for WC1.

**Figure 1 B – source data WC2:** Western blots were used to detect expression of WC2 in the indicated samples for Figure 1 B.

**Figure 1 B – source data WC2_labeled:** Figure with the area highlighted was used to develop the Figure 1 B for WC2.

**Figure 1 B – source data RGB1:** Western blots were used to detect expression of RGB1 in the indicated samples for Figure 1 B.

**Figure 1 B – source data RGB1_labeled:** Figure with the area highlighted was used to develop the Figure 1 B for RGB1.

**Figure 1 B – source data LC:** Ponceau S staining was used to detect loading control of the indicated samples for Figure 1 B.

**Figure 1 B – source data LC_labeled:** Figure with the area highlighted was used to develop the Figure 1 B for LC.

### Source data for Figure 1 F

**Figure 1 F – source data FRQ_s:** Western blots were used to detect expression of FRQ in the indicated samples for Figure 1 F. (s: short exposure)

**Figure 1 F – source data FRQ_s_labeled:** Figure with the area highlighted was used to develop the Figure 1 F for FRQ. (s: short exposure)

**Figure 1 F – source data FRQ_l:** Western blots were used to detect expression of FRQ in the indicated samples for Figure 1 F. (l: long exposure)

**Figure 1 F – source data FRQ_l_labeled:** Figure with the area highlighted was used to develop the Figure 1 F for FRQ. (l: long exposure)

**Figure 1 F – source data WC1_s:** Western blots were used to detect expression of WC1 in the indicated samples for Figure 1 F. (s: short exposure)

**Figure 1 F – source data WC1_s_labeled:** Figure with the area highlighted was used to develop the Figure 1 F for WC1. (s: short exposure)

**Figure 1 F – source data WC1_l:** Western blots were used to detect expression of WC1 in the indicated samples for Figure 1 F. (l: long exposure)

**Figure 1 F – source data WC1_l_labeled:** Figure with the area highlighted was used to develop the Figure 1 F for WC1. (l: long exposure)

**Figure 1 F – source data WC2_s:** Western blots were used to detect expression of WC2 in the indicated samples for Figure 1 F. (s: short exposure)

**Figure 1 F – source data WC2_s_labeled:** Figure with the area highlighted was used to develop the Figure 1 F for WC2. (s: short exposure)

**Figure 1 F – source data WC2_l:** Western blots were used to detect expression of WC2 in the indicated samples for Figure 1 F. (l: long exposure)

**Figure 1 F – source data WC2_l_labeled:** Figure with the area highlighted was used to develop the Figure 1 F for WC2. (l: long exposure)

**Figure 1 F – source data LC:** Ponceau S staining was used to detect loading control of the indicated samples for Figure 1 F.

**Figure 1 F – source data LC_labeled:** Figure with the area highlighted was used to develop the Figure 1 F for LC.

### Source data for Figure 2 A

**Figure 2 A - source data_001glc:** Western blots were used to detect expression of FRQ in the indicated samples grown in 0.01% glucose containing medium for Figure 2 A.

**Figure 2 A - source data_001glc_labeled:** Figure with the area highlighted was used to develop the Figure 2 A for FRQ in 0.01% glucose.

**Figure 2 A - source data_2glc:** Western blots were used to detect expression of FRQ in the indicated samples grown in 2% glucose containing medium for Figure 2 A.

**Figure 2 A - source data_2glc_labeled:** Figure with the area highlighted was used to develop the Figure 2 A for FRQ in 2% glucose.

**Figure 2 A - source data_LC_001glc:** Ponceau S staining was used to detect loading control of the indicated samples for Figure 2 A. (0.01% glucose)

**Figure 2 A - source data_LC_001glc_labeled:** Figure with the area highlighted was used to develop the Figure 2 A for LC. (0.01% glucose)

**Figure 2 A - source data_LC_2glc:** Ponceau S staining was used to detect loading control of the indicated samples for Figure 2 A. (2% glucose)

**Figure 2 A - source data_LC_2glc_labeled:** Figure with the area highlighted was used to develop the Figure 2 A for LC. (2% glucose)

### Source data for Figure 2 B

**Figure 2 B – source data_FRQ:** Western blots were used to detect expression of FRQ in the indicated samples for Figure 2 B.

**Figure 2 B – source data_FRQ_labeled:** Figure with the area highlighted was used to develop the Figure 2 B for FRQ.

**Figure 2 B – source data_WC1:** Western blots were used to detect expression of WC1 in the indicated samples for Figure 2 B.

**Figure 2 B – source data_WC1_labeled:** Figure with the area highlighted was used to develop the Figure 2 B for WC1.

**Figure 2 B – source data_WC2:** Western blots were used to detect expression of WC1 in the indicated samples for Figure 2 B.

**Figure 2 B – source data_WC2_labeled:** Figure with the area highlighted was used to develop the Figure 2 B for WC2.

**Figure 2 B - source data_LC:** Ponceau S staining was used to detect loading control of the indicated samples for Figure 2 B.

**Figure 2 B - source data_LC_labeled:** Figure with the area highlighted was used to develop the Figure 2 B for LC.

### Source data for Figure 3 A

**Figure 3 A – source data WC1_s:** Western blots were used to detect expression of WC1 in the indicated samples for Figure 3 A. (s: short exposure)

**Figure 3 A – source data WC1_s_labeled:** Figure with the area highlighted was used to develop the Figure 3 A for WC1. (s: short exposure)

**Figure 3 A – source data WC1_l:** Western blots were used to detect expression of WC1 in the indicated samples for Figure 3 A. (l: long exposure)

**Figure 3 A – source data WC1_l_labeled:** Figure with the area highlighted was used to develop the Figure 3 A for WC1. (l: long exposure)

**Figure 3 A – source data WC2_s:** Western blots were used to detect expression of WC2 in the indicated samples for Figure 3 A. (s: short exposure)

**Figure 3 A – source data WC2_s_labeled:** Figure with the area highlighted was used to develop the Figure 3 A for WC2. (s: short exposure)

**Figure 3 A – source data WC2_l:** Western blots were used to detect expression of WC2 in the indicated samples for Figure 3 A. (l: long exposure)

**Figure 3 A – source data WC2_l_labeled:** Figure with the area highlighted was used to develop the Figure 3 A for WC2. (l: long exposure)

**Figure 3 A – source data LC:** Ponceau S staining was used to detect loading control of the indicated samples for Figure 3 A.

**Figure 3 A – source data LC_ for WC1 labeled:** Figure with the area highlighted was used to develop the Figure 3 A for LC for WC1.

**Figure 3 A – source data LC_ for WC2 labeled:** Figure with the area highlighted was used to develop the Figure 3 A for LC for WC2.

### Source data for Figure 3 C

**Figure 3 C – source data_FRQ:** Western blots were used to detect expression of FRQ in the indicated samples for Figure 3 C.

**Figure 3 C – source data_FRQ_labeled:** Figure with the area highlighted was used to develop the Figure 3 C for FRQ.

**Figure 3 C – source data_WC1:** Western blots were used to detect expression of WC1 in the indicated samples for Figure 3 C.

**Figure 3 C – source data_WC1_labeled:** Figure with the area highlighted was used to develop the Figure 3 C for WC1.

**Figure 3 C – source data_WC2:** Western blots were used to detect expression of WC2 in the indicated samples for Figure 3 C.

**Figure 3 C – source data_WC2_labeled:** Figure with the area highlighted was used to develop the Figure 3 C for WC2.

**Figure 3 C – source data LC:** Ponceau S staining was used to detect loading control of the indicated samples for Figure 3 C.

**Figure 3 C – source data LC_ for FRQ_WC2 labeled:** Figure with the area highlighted was used to develop the Figure 3 C for LC for FRQ and WC2.

**Figure 3 C – source data LC_ for WC1 labeled:** Figure with the area highlighted was used to develop the Figure 3 C for LC for WC1.

### Source data for Figure 3 D

**Figure 3 D – source data FRQ_s:** Western blots were used to detect expression of FRQ in the indicated samples for Figure 3 D. (s: short exposure)

**Figure 3 D – source data FRQ_s_labeled:** Figure with the area highlighted was used to develop the Figure 3 D for FRQ. (s: short exposure)

**Figure 3 D – source data FRQ_l:** Western blots were used to detect expression of FRQ in the indicated samples for Figure 3 D. (l: long exposure)

**Figure 3 D – source data FRQ_l_labeled:** Figure with the area highlighted was used to develop the Figure 3 D for FRQ. (l: long exposure)

**Figure 3 D – source data_WC1:** Western blots were used to detect expression of WC1 in the indicated samples for Figure 3 D.

**Figure 3 D – source data_WC1_labeled:** Figure with the area highlighted was used to develop the Figure 3 D for WC1.

**Figure 3 D – source data_WC2:** Western blots were used to detect expression of WC2 in the indicated samples for Figure 3 D.

**Figure 3 D – source data_WC2_labeled:** Figure with the area highlighted was used to develop the Figure 3 D for WC2.

**Figure 3 D – source data LC:** Ponceau S staining was used to detect loading control of the indicated samples for Figure 3 D.

**Figure 3 D – source data LC_ for FRQ labeled:** Figure with the area highlighted was used to develop the Figure 3 D for LC for FRQ.

**Figure 3 D – source data LC_ for WC1 and WC2 labeled:** Figure with the area highlighted was used to develop the Figure 3 D for LC for WC1 and for WC2.

### Source data for Figure 3 E

**Figure 3 E – source data FRQ:** Western blots were used to detect expression of FRQ in the indicated samples for Figure 3 E.

**Figure 3 E – source data FRQ_labeled:** Figure with the area highlighted was used to develop the Figure 3 E for FRQ.

**Figure 3 E – source data WC1:** Western blots were used to detect expression of WC1 in the indicated samples for Figure 3 E.

**Figure 3 E – source data WC1_labeled:** Figure with the area highlighted was used to develop the Figure 3 E for WC1.

**Figure 3 E – source data WC2:** Western blots were used to detect expression of WC2 in the indicated samples for Figure 3 E.

**Figure 3 E – source data WC2_labeled:** Figure with the area highlighted was used to develop the Figure 3 E for WC2.

**Figure 3 E – source data LC:** Ponceau S staining was used to detect loading control of the indicated samples for Figure 3 E.

**Figure 3 E – source data LC_labeled:** Figure with the area highlighted was used to develop the Figure 3 E for LC.

### Source data for Figure 3 G

**Figure 3 G – source data FRQ:** Western blots were used to detect expression of FRQ in the indicated samples for Figure 3 G.

**Figure 3 G – source data FRQ_labeled:** Figure with the area highlighted was used to develop the Figure 3 G for FRQ.

**Figure 3 G – source data WC1:** Western blots were used to detect expression of WC1 in the indicated samples for Figure 3 G.

**Figure 3 G – source data WC1_labeled:** Figure with the area highlighted was used to develop the Figure 3 G for WC1.

**Figure 3 G – source data WC2:** Western blots were used to detect expression of WC2 in the indicated samples for Figure 3 G.

**Figure 3 G – source data WC2_labeled:** Figure with the area highlighted was used to develop the Figure 3 G for WC2.

**Figure 3 G – source data LC for FRQ:** Ponceau S staining was used to detect loading control of the indicated samples for Figure 3 G for FRQ.

**Figure 3 G – source data LC for FRQ_labeled:** Figure with the area highlighted was used to develop the Figure 3 G for LC for FRQ.

**Figure 3 G – source data LC for WC1_labeled:** Figure with the area highlighted was used to develop the Figure 3 G for LC for WC1.

**Figure 3 G – source data LC for WC2_labeled:** Figure with the area highlighted was used to develop the Figure 3 G for LC for WC2.

### Source data for Supplemental Figure 1 B

**Supplemental Figure 1 B – source data FRQ:** Western blots were used to detect expression of FRQ in the indicated samples for Supplemental Figure 1 B.

**Supplemental Figure 1 B – source data FRQ_labeled:** Figure with the area highlighted was used to develop the Supplemental Figure 1 B for FRQ.

**Supplemental Figure 1 B – source data WC1:** Western blots were used to detect expression of WC1 in the indicated samples for Supplemental Figure 1 B.

**Supplemental Figure 1 B – source data WC1_labeled:** Figure with the area highlighted was used to develop the Supplemental Figure1 B for WC1.

**Supplemental Figure 1 B – source data WC2:** Western blots were used to detect expression of WC2 in the indicated samples for Supplemental Figure 1 B.

**Supplemental Figure 1 B – source data WC2_labeled:** Figure with the area highlighted was used to develop the Supplemental Figure 1 B for WC2.

**Supplemental Figure 1 B – source data LC:** Ponceau S staining was used to detect loading control of the indicated samples for Supplemental Figure 1 B for FRQ.

**Supplemental Figure 1 B – source data LC _labeled:** Figure with the area highlighted was used to develop the Supplemental Figure 1 B for LC.

## Notes

### Competing Interest Statement

The authors have declared no competing interest.

## References

1. Adhvaryu, K., Firoozi, G., Motavaze, K., and Lakin-Thomas, P. (2016). PRD-1, a Component of the Circadian System of Neurospora crassa, Is a Member of the DEAD-box RNA Helicase Family. J Biol Rhythms 31, 258–271.

2. Anders, S., Pyl, P.T., and Huber, W. (2015). HTSeq-a Python framework to work with high-throughput sequencing data. Bioinformatics 31, 166–169.

3. Andrews, S. (2010). A Quality Control Tool for High Throughput Sequence Data. FastQC.

4. Aronson, B.D., Johnson, K.A., and Dunlap, J.C. (1994). Circadian clock locus frequency: protein encoded by a single open reading frame defines period length and temperature compensation. Proc Natl Acad Sci U S A 91, 7683–7687.

5. Ashe, M.P., De Long, S.K., and Sachs, A.B. (2000). Glucose depletion rapidly inhibits translation initiation in yeast. Mol Biol Cell 11, 833–848.

6. Baron, K.G., and Reid, K.J. (2014). Circadian misalignment and health. Int Rev Psychiatry 26, 139–154.

7. Barraza, C.E., Solari, C.A., Marcovich, I., Kershaw, C., Galello, F., Rossi, S., Ashe, M.P., and Portela, P. (2017). The role of PKA in the translational response to heat stress in Saccharomyces cerevisiae. PLoS One 12, e0185416.

8. Bell-Pedersen, D., Dunlap, J.C., and Loros, J.J. (1992). The Neurospora circadian clock-controlled gene, ccg-2, is allelic to eas and encodes a fungal hydrophobin required for formation of the conidial rodlet layer. Genes Dev 6, 2382–2394.

9. Bell-Pedersen, D., Dunlap, J.C., and Loros, J.J. (1996). Distinct cis-acting elements mediate clock, light, and developmental regulation of the Neurospora crassa eas (ccg-2) gene. Mol Cell Biol 16, 513–521.

10. Benocci, T., Aguilar-Pontes, M.V., Zhou, M., Seiboth, B., and de Vries, R.P. (2017). Regulators of plant biomass degradation in ascomycetous fungi. Biotechnol Biofuels 10, 152.

11. Castermans, D., Somers, I., Kriel, J., Louwet, W., Wera, S., Versele, M., Janssens, V., and Thevelein, J.M. (2012). Glucose-induced posttranslational activation of protein phosphatases PP2A and PP1 in yeast. Cell Res 22, 1058–1077.

12. Cheng, P., He, Q., He, Q., Wang, L., and Liu, Y. (2005). Regulation of the Neurospora circadian clock by an RNA helicase. Genes Dev 19, 234–241.

13. Cheng, P., Yang, Y., and Liu, Y. (2001). Interlocked feedback loops contribute to the robustness of the Neurospora circadian clock. Proc Natl Acad Sci U S A 98, 7408–7413.

14. Colot, H.V., Park, G., Turner, G.E., Ringelberg, C., Crew, C.M., Litvinkova, L., Weiss, R.L., Borkovich, K.A., and Dunlap, J.C. (2006). A high-throughput gene knockout procedure for Neurospora reveals functions for multiple transcription factors. Proc Natl Acad Sci U S A 103, 10352–10357.

15. Conrad, M., Schothorst, J., Kankipati, H.N., Van Zeebroeck, G., Rubio-Texeira, M., and Thevelein, J.M. (2014). Nutrient sensing and signaling in the yeast Saccharomyces cerevisiae. Fems Microbiology Reviews 38, 254–299.

16. Corral-Ramos, C., Barrios, R., Ayte, J., and Hidalgo, E. (2021). TOR and MAP kinase pathways synergistically regulate autophagy in response to nutrient depletion in fission yeast. Autophagy, 1-16.

17. Cusick, K.D., Fitzgerald, L.A., Pirlo, R.K., Cockrell, A.L., Petersen, E.R., and Biffinger, J.C. (2014). Selection and Evaluation of Reference Genes for Expression Studies with Quantitative PCR in the Model Fungus Neurospora crassa under Different Environmental Conditions in Continuous Culture. Plos One 9.

18. Dobin, A., Davis, C.A., Schlesinger, F., Drenkow, J., Zaleski, C., Jha, S., Batut, P., Chaisson, M., and Gingeras, T.R. (2013). STAR: ultrafast universal RNA-seq aligner. Bioinformatics 29, 15–21.

19. Emerson, J.M., Bartholomai, B.M., Ringelberg, C.S., Baker, S.E., Loros, J.J., and Dunlap, J.C. (2015). period-1 encodes an ATP-dependent RNA helicase that influences nutritional compensation of the Neurospora circadian clock. Proc Natl Acad Sci U S A 112, 15707–15712.

20. Froehlich, A.C., Liu, Y., Loros, J.J., and Dunlap, J.C. (2002). White Collar-1, a circadian blue light photoreceptor, binding to the frequency promoter. Science 297, 815–819.

21. Gorl, M., Merrow, M., Huttner, B., Johnson, J., Roenneberg, T., and Brunner, M. (2001). A PEST-like element in FREQUENCY determines the length of the circadian period in Neurospora crassa. EMBO J 20, 7074–7084.

22. Gyongyosi, N., and Kaldi, K. (2014). Interconnections of reactive oxygen species homeostasis and circadian rhythm in Neurospora crassa. Antioxid Redox Signal 20, 3007–3023.

23. Gyongyosi, N., Nagy, D., Makara, K., Ella, K., and Kaldi, K. (2013). Reactive oxygen species can modulate circadian phase and period in Neurospora crassa. Free Radic Biol Med 58, 134–143.

24. Gyongyosi, N., Szoke, A., Ella, K., and Kaldi, K. (2017). The small G protein RAS2 is involved in the metabolic compensation of the circadian clock in the circadian model Neurospora crassa. J Biol Chem 292, 14929–14939.

25. Hallett, J.E.H., Luo, X.X., and Capaldi, A.P. (2014). State Transitions in the TORC1 Signaling Pathway and Information Processing in Saccharomyces cerevisiae. Genetics 198, 773–U452.

26. He, Q., Cha, J., He, Q., Lee, H.C., Yang, Y., and Liu, Y. (2006). CKI and CKII mediate the FREQUENCY-dependent phosphorylation of the WHITE COLLAR complex to close the Neurospora circadian negative feedback loop. Genes Dev 20, 2552–2565.

27. He, Q.Y., Cheng, P., Yang, Y.H., Wang, L.X., Gardner, K.H., and Liu, Y. (2002). White collar-1, a DNA binding transcription factor and a light sensor. Science 297, 840–843.

28. Huang, G., Chen, S., Li, S., Cha, J., Long, C., Li, L., He, Q., and Liu, Y. (2007). Protein kinase A and casein kinases mediate sequential phosphorylation events in the circadian negative feedback loop. Genes Dev 21, 3283–3295.

29. Hurley, J.H., Dasgupta, A., Andrews, P., Crowell, A.M., Ringelberg, C., Loros, J.J., and Dunlap, J.C. (2015). A Tool Set for the Genome-Wide Analysis of Neurospora crassa by RT-PCR. G3-Genes Genomes Genetics 5, 2043-2049.

30. Hurley, J.M., Dasgupta, A., Emerson, J.M., Zhou, X., Ringelberg, C.S., Knabe, N., Lipzen, A.M., Lindquist, E.A., Daum, C.G., Barry, K.W., et al. (2014). Analysis of clock-regulated genes in Neurospora reveals widespread posttranscriptional control of metabolic potential. Proc Natl Acad Sci U S A 111, 16995–17002.

31. Jona, G., Choder, M., and Gileadi, O. (2000). Glucose starvation induces a drastic reduction in the rates of both transcription and degradation of mRNA in yeast. Biochim Biophys Acta 1491, 37–48.

32. Kaldenhoff, R., and Russo, V.E. (1993). Promoter analysis of the bli-7/eas gene. Curr Genet 24, 394–399.

33. Kodadek, T., Sikder, D., and Nalley, K. (2006). Keeping transcriptional activators under control. Cell 127, 261–264.

34. Larrondo, L.F., Olivares-Yanez, C., Baker, C.L., Loros, J.J., and Dunlap, J.C. (2015). Circadian rhythms. Decoupling circadian clock protein turnover from circadian period determination. Science 347, 1257277.

35. Lee, H.Y., Itahana, Y., Schuechner, S., Fukuda, M., Je, H.S., Ogris, E., Virshup, D.M., and Itahana, K. (2018). Ca(2+)-dependent demethylation of phosphatase PP2Ac promotes glucose deprivation-induced cell death independently of inhibiting glycolysis. Sci Signal 11.

36. Lee, K., Loros, J.J., and Dunlap, J.C. (2000). Interconnected feedback loops in the Neurospora circadian system. Science 289, 107–110.

37. Leipheimer, J., Bloom, A.L.M., and Panepinto, J.C. (2019). Protein Kinases at the Intersection of Translation and Virulence. Front Cell Infect Microbiol 9, 318.

38. Li, H., Handsaker, B., Wysoker, A., Fennell, T., Ruan, J., Homer, N., Marth, G., Abecasis, G., Durbin, R., and Genome Project Data Processing, S. (2009). The Sequence Alignment/Map format and SAMtools. Bioinformatics 25, 2078-2079.

39. Li, L., and Borkovich, K.A. (2006a). GPR-4 is a predicted G-protein-coupled receptor required for carbon source-dependent asexual growth and development in Neurospora crassa. Eukaryot Cell 5, 1287–1300.

40. Li, L.D., and Borkovich, K.A. (2006b). GPR-4 is a predicted G-protein-coupled receptor required for carbon source-dependent asexual growth and development in Neurospora crassa. Eukaryotic Cell 5, 1287–1300.

41. Liu, X., Chen, A., Caicedo-Casso, A., Cui, G., Du, M., He, Q., Lim, S., Kim, H.J., Hong, C.I., and Liu, Y. (2019). FRQ-CK1 interaction determines the period of circadian rhythms in Neurospora. Nat Commun 10, 4352.

42. Llanos, A., Francois, J.M., and Parrou, J.L. (2015). Tracking the best reference genes for RT-qPCR data normalization in filamentous fungi. Bmc Genomics 16.

43. Love, M.I., Huber, W., and Anders, S. (2014). Moderated estimation of fold change and dispersion for RNA-seq data with DESeq2. Genome Biol 15.

44. Luo, C.H., Loros, J.J., and Dunlap, J.C. (1998). Nuclear localization is required for function of the essential clock protein FRQ. Embo Journal 17, 1228–1235.

45. Malzahn, E., Ciprianidis, S., Kaldi, K., Schafmeier, T., and Brunner, M. (2010). Photoadaptation in Neurospora by competitive interaction of activating and inhibitory LOV domains. Cell 142, 762–772.

46. Margolin, B., S., Freitag, M., Selker, E. U. (1997). Improved plasmids for gene targeting at the his-3 locus of Neurospora crassa by electroporation. Fungal Genetics Reports 44.

47. McCluskey, K. (2003). The Fungal Genetics Stock Center: from molds to molecules. Adv Appl Microbiol 52, 245–262.

48. Mi, H.Y., Muruganujan, A., Casagrande, J.T., and Thomas, P.D. (2013). Large-scale gene function analysis with the PANTHER classification system. Nat Protoc 8, 1551–1566.

49. Nitsche, B.M., Jorgensen, T.R., Akeroyd, M., Meyer, V., and Ram, A.F. (2012). The carbon starvation response of Aspergillus niger during submerged cultivation: insights from the transcriptome and secretome. BMC Genomics 13, 380.

50. Nosanchuk, J.D., Stark, R.E., and Casadevall, A. (2015). Fungal Melanin: What do We Know About Structure? Front Microbiol 6, 1463.

51. Olivares-Yanez, C., Emerson, J., Kettenbach, A., Loros, J.J., Dunlap, J.C., and Larrondo, L.F. (2016). Modulation of Circadian Gene Expression and Metabolic Compensation by the RCO-1 Corepressor of Neurospora crassa. Genetics 204, 163–176.

52. Punga, T., Bengoechea-Alonso, M.T., and Ericsson, J. (2006). Phosphorylation and ubiquitination of the transcription factor sterol regulatory element-binding protein-1 in response to DNA binding. Journal of Biological Chemistry 281, 25278–25286.

53. Quan, Z., Cao, L., Tang, Y., Yan, Y., Oliver, S.G., and Zhang, N. (2015). The Yeast GSK-3 Homologue Mck1 Is a Key Controller of Quiescence Entry and Chronological Lifespan. PLoS Genet 11, e1005282.

54. Querfurth, C., Diernfellner, A.C., Gin, E., Malzahn, E., Hofer, T., and Brunner, M. (2011). Circadian conformational change of the Neurospora clock protein FREQUENCY triggered by clustered hyperphosphorylation of a basic domain. Mol Cell 43, 713–722.

55. R Core Team (2020). R: A language and environment for statistical computing. R Foundation for Statistical Computing.

56. Sancar, C., Sancar, G., Ha, N., Cesbron, F., and Brunner, M. (2015). Dawn- and dusk-phased circadian transcription rhythms coordinate anabolic and catabolic functions in Neurospora. BMC Biol 13, 17.

57. Sancar, G., Sancar, C., and Brunner, M. (2012). Metabolic compensation of the Neurospora clock by a glucose-dependent feedback of the circadian repressor CSP1 on the core oscillator. Genes Dev 26, 2435–2442.

58. Schafmeier, T., Diernfellner, A., Schafer, A., Dintsis, O., Neiss, A., and Brunner, M. (2008). Circadian activity and abundance rhythms of the Neurospora clock transcription factor WCC associated with rapid nucleo-cytoplasmic shuttling. Gene Dev 22, 3397–3402.

59. Schafmeier, T., Haase, A., Kaldi, K., Scholz, J., Fuchs, M., and Brunner, M. (2005). Transcriptional feedback of Neurospora circadian clock gene by phosphorylation-dependent inactivation of its transcription factor. Cell 122, 235–246.

60. Schafmeier, T., Kaldi, K., Diernfellner, A., Mohr, C., and Brunner, M. (2006). Phosphorylation-dependent maturation of Neurospora circadian clock protein from a nuclear repressor toward a cytoplasmic activator. Genes Dev 20, 297–306.

61. Sokolovsky, V.Y., Lauter, F.R., Mullerrober, B., Ricci, M., Schmidhauser, T.J., and Russo, V.E.A. (1992). Nitrogen Regulation of Blue Light-Inducible Genes in Neurospora-Crassa. Journal of General Microbiology 138, 2045–2049.

62. Springer, M.L. (1993). Genetic control of fungal differentiation: the three sporulation pathways of Neurospora crassa. Bioessays 15, 365–374.

63. Tataroglu, O., Lauinger, L., Sancar, G., Jakob, K., Brunner, M., and Diernfellner, A.C. (2012). Glycogen synthase kinase is a regulator of the circadian clock of Neurospora crassa. J Biol Chem 287, 36936–36943.

64. Vogel, H.J. (1964). Distribution of lysine pathways among fungi: evolutionary implications. Am. Naturalist, 435-446.

65. Wang, B., Li, J., Gao, J., Cai, P., Han, X., and Tian, C. (2017). Identification and characterization of the glucose dual-affinity transport system in Neurospora crassa: pleiotropic roles in nutrient transport, signaling, and carbon catabolite repression. Biotechnol Biofuels 10, 17.

66. Yang, Y., He, Q., Cheng, P., Wrage, P., Yarden, O., and Liu, Y. (2004). Distinct roles for PP1 and PP2A in the Neurospora circadian clock. Genes Dev 18, 255–260.

67. Yates, A.D., Achuthan, P., Akanni, W., Allen, J., Allen, J., Alvarez-Jarreta, J., Amode, M.R., Armean, I.M., Azov, A.G., Bennett, R., et al. (2020). Ensembl 2020. Nucleic Acids Res 48, D682–D688.

